# Sustained multigenerational fitness benefits of natural immigration

**DOI:** 10.64898/2026.05.13.724961

**Authors:** Jeremy Summers, Elissa J. Cosgrove, Tori Bakley, Sahas Barve, Reed Bowman, John W. Fitzpatrick, Nancy Chen

## Abstract

The fitness of immigrants and their descendants determines the effectiveness of gene flow. Genetic incompatibilities or outbreeding depression can limit the spread of novel alleles, while highly fit immigrant lineages can hasten introgression. These fitness effects of gene flow can also differ between generations as immigrant and resident haplotypes recombine. Understanding the genetic factors that shape immigrant fitness over multiple generations is increasingly important as habitat fragmentation threatens populations by reducing genetic variation and leading to increased levels of inbreeding. Few studies have measured the multigenerational fitness of immigrant lineages, especially within populations that had histories of high gene flow. We used 33 years of life history and pedigree data on a population of Florida scrub-jays (*Aphelocoma coerulescens*) with historically high immigration to quantify the fitness of immigrants and their descendants. We compared the fitness of immigrants and residents as well as their resulting descendants (F1, F2, etc.) to determine the composite genetic effects responsible for fitness differences. We found evidence of additive benefits of immigrant ancestry and heterosis driven by non-additive effects that persists for multiple generations. These results are promising for conservation efforts aiming to increase connectivity and illustrate the complex dynamics that determine the rates of introgression in natural populations.

## Introduction

Habitat fragmentation is isolating natural populations globally (Templeton et al. 1990; Simberloff 1999; Wilson et al. 2016; Fletcher et al. 2018; Exposito-Alonso et al. 2022), reshaping their evolutionary trajectories and contributing to threats of extinction. Dispersal and subsequent gene flow between habitat patches may play an increasingly important role in governing the dynamics and persistence of natural populations (Hedrick and Garcia-Dorado 2016; Forester et al. 2022; Grummer et al. 2022). The consequences of gene flow are determined by the fitness of dispersing individuals and their many types of descendants over generations (Roff and Emerson 2006; Dickel et al. 2023; Goedert et al. 2025). Thus, the multigenerational fitness effects of gene flow can be complex, and characterizing these effects is vital for understanding contemporary evolution and guiding conservation efforts. To date, very few studies have quantified multigenerational fitness outcomes of natural immigration in the wild (Martinig et al. 2020; Dickel et al. 2023; Goedert et al. 2025; Saatoglu et al. 2025).

The fitness of immigrants and their descendants depends on the genetic architecture of fitness as well as genetic and ecological similarities between immigrant source and resident populations (Goedert et al. 2025). The fitness of incoming immigrants can be lower than that of residents if there is strong local adaptation, but higher if they originate from source populations with less inbreeding depression or if they bring in new beneficial genetic variation (Lenormand 2002; Bell et al. 2019).

Offspring of immigrant - resident pairings (F1s) could have higher heterozygosity, masking recessive deleterious alleles from either population and resulting in heterosis (Charlesworth and Charlesworth 1987; Lynch 1991; Frankham 2015). On the other hand, descendants of immigrants could have decreased fitness due to incompatibilities between immigrant and resident alleles or the disruption of co-adapted gene complexes through recombination between immigrant and resident genomes (Lenormand 2002).

The relative fitness of residents, immigrants, and their various types of descendants (F1, F2, resident backcross, immigrant backcross, etc.) reflect the net outcomes of these different effects, which may also vary between sexes and among fitness components (Roff and Emerson 2006).

Comparing the fitness of individuals across different immigrant ancestry categories can provide insights on the genetic architecture of fitness. A powerful framework for inferring complex genetic architectures is line-cross theory, which leverages phenotypic means from known crosses between populations to estimate the net composite genetic effect of all additive, dominant, and epistatic loci that influence a trait (Mather and Jinks 1978; Hill 1982). Line-cross theory can also be used to generate a series of specific comparisons to test for evidence of non-additive effects (Mather and Jinks 1978; Hill 1982; Lynch 1991), and has been frequently applied in agricultural and experimental settings with the ability to generate controlled crosses, revealing the role of non-additive effects in both reproductive isolation and heterosis (Yu et al. 1997; Costa e Silva et al. 2012; Owusu et al. 2022; Le Rouzic et al. 2024), and recently was applied to a natural population of song sparrows to study the fitness effects of immigration (Dickel et al. 2023). A better understanding of the additive, dominance, and epistatic genetic effects underlying fitness differences between individuals of varying immigrant ancestry is needed to better predict the outcomes of introgression and evolutionary dynamics in spatially structured populations.

Theoretical and empirical studies suggest that the fitness outcomes of immigration are likely to be influenced by non-additive genetic effects and be context-specific (Dickel et al. 2023; Reid et al. 2024; Goedert et al. 2025). Non-additive genetic effects are expected to be larger in life history traits compared to morphological traits in natural populations due to lower additive genetic variation for fitness in populations near fitness optima (Roff and Emerson 2006; Burch et al. 2024). However, theory also predicts that simpler genetic architectures involving additive and dominance effects are more likely among less diverged populations, while epistatic effects become more common among more diverged populations (Burch et al. 2024). The success of genetic rescue efforts (Frankham 2015; Whiteley et al. 2015; Fitzpatrick et al. 2020) suggests that dominant genetic effects may play a large role in the fitness consequences of immigration, especially in small, inbred populations with relatively low levels of divergence. One of very few studies to date that quantified the multigenerational fitness outcomes of immigrants and their descendants found evidence of strong non-additive effects of immigrant ancestry within an island population of song sparrows (*Melospiza melodia*) with relatively low levels of natural immigration (Dickel et al. 2023). This study and others have documented sex-specific fitness effects of immigration (Martinig et al. 2020; Reid et al. 2024), consistent with Haldane’s rule for interspecific crosses (Orr 1997). Given the complexity of fitness outcomes of immigration, more empirical studies that quantify the fitness of multiple generations of immigrant descendants are needed, particularly in populations with varying demographic histories.

Here, we quantified the fitness of immigrants and their descendants across generations and applied line-cross theory to infer the genetic architecture of fitness in a population of cooperatively breeding Florida scrub-jays (*Aphelocoma coerulescens*) that has experienced rapid declines in immigration from historically high levels (Chen et al. 2016; Summers et al. 2024). The Florida scrub-jay is a habitat specialist that is restricted to the xeric oak scrub of Florida and is threatened by range-wide habitat loss and fragmentation (Woolfenden and Fitzpatrick 1984; Boughton and Bowman 2011).

The population at Archbold Biological Station (hereafter ABS) has been extensively monitored since 1969, accruing direct measurements of immigration and extensive pedigree data (Woolfenden and Fitzpatrick 1984). Previous work showed that temporal declines in immigration have contributed to increased inbreeding and decreased fitness due to inbreeding depression (Chen et al. 2016). We first compared the fitness of different immigrant ancestry groups to test for non-additive effects, then used line-cross analysis to determine the specific composite genetic effects that underlie the fitness differences observed in our comparisons. Given observed inbreeding depression, we expect non-additive effects to result in heterosis within the Florida scrub-jays at ABS, but otherwise we do not expect to see evidence of outbreeding depression due to historic and ongoing gene flow. This work leverages more than 30 years of complete pedigree data to elucidate the short and long term consequences of gene flow in a population experiencing increasing isolation.

## Methods and Materials

### Study species and system

The population of Florida scrub-jays at ABS is among the largest within the Lake Wales Ridge metapopulation (Coulon et al. 2008) and receives substantial immigration from the smaller, surrounding populations (Boughton and Bowman 2011; Chen et al. 2016; Summers et al. 2024). Researchers at ABS have monitored the population since 1969, uniquely banding all individuals within the study tract at 11 days of age, or upon discovery for immigrants (Woolfenden and Fitzpatrick 1984). Florida scrub-jays are highly philopatric and remain within well-defined territories year-round, facilitating tracking individuals across their lifetimes (Woolfenden and Fitzpatrick 1984) and resulting in a 16-generation population pedigree (Chen et al. 2016). Emigration rates are low from our study site, allowing us to infer effective mortality in particular for juveniles and breeders, who are far less likely to disperse than adult non-breeding helpers (Woolfenden and Fitzpatrick 1984). Individuals are sexed via field methods (Woolfenden and Fitzpatrick 1984) or molecular sexing starting in 1999 (Woolfenden and Fitzpatrick 2020). All work was approved by the Cornell University Institutional Animal Care and Use Committee (IACUC 2010-0015) and authorized by permits from the US Fish and Wildlife Service (TE824723-8), the US Geological Survey (banding permit 07732), and the Florida Fish and Wildlife Conservation Commission (LSSC-10-00205).

### Ancestry categories

Due to expansions in the study area in the 1980s, we assigned all individuals present on the study tract in 1986 as residents and all subsequent individuals born outside our study tract as immigrants. We then assigned all other individuals to ancestry categories based on the number of immigrant grandparents they had: zero for residents (R), one for resident backcrosses (Rbc), two for F1s or F2s (F2s are offspring of two F1 individuals), three for immigrant backcrosses (Ibc), and four for immigrant founders (I). Individuals born on the study tract with four immigrant grandparents were assigned as immigrant offspring (IO) and considered immigrants when quantifying immigrant ancestry of their descendants. Individuals with unknown grandparents were assigned a separate category excluded from our fitness comparisons and line cross analysis, but were included in our models for the purpose of fitting random effects (*N* = 105). For all comparisons and our line-cross analysis, we used immigrant offspring as our immigrant value to avoid including additional environmental variation from individuals hatching in different locations and dispersing to the study tract. See Supplementary Figure S1 for the distribution of ancestry categories across cohorts.

### Fitness components

We investigated the effect of immigrant ancestry on a set of fitness metrics that combined account for full lifetime reproductive success. For adults who successfully become a breeder, we investigated their breeder lifetime reproductive success (LRS) and its sub-components: annual breeder survival and annual reproductive success (ARS). For LRS, we quantified the total number of banded (i.e., 11-day-old) juveniles produced over the lifetimes of each individual. We only included individuals born between 1991 (the first cohort with F1 individuals who became breeders) and 2014 (the last cohort with <5% of individuals still alive; the 11 surviving individuals are included; *N =* 703 individuals). We defined breeder survival as survival from the start of one breeding season in March (Woolfenden and Fitzpatrick 1984) to the start of the next breeding season, and we measured ARS as the number of 11-day-old juveniles produced by an individual within one year. For the rare cases of multiple broods in a single year, we removed nests that failed to produce any fledglings (18-day-old individuals) (*N* = 163 nests) to avoid inflating ARS for pairs that attempted to breed again after nest failure. Breeder survival and ARS analyses included individuals born between 1993 (the first year F1 individuals bred) and 2023 (*N* = 3,700 annual records). All breeder analyses included both locally hatched and immigrant founder individuals. As Florida scrub-jays start dispersing after their 1st year, we assigned immigrant founders to cohorts based on their minimum likely age (Woolfenden and Fitzpatrick 1984; Suh et al. 2020).

To investigate the effect of immigrant ancestry on recruitment, we estimated juvenile survival and subsequent likelihood to establish as a breeder (breeder transition). We defined juvenile survival from 11 days to the start of the next breeding season (March) of the next year. We restricted this analysis to cohorts where >95% of individuals were sexed molecularly (cohorts 1999-2013; *N* = 3,192 individuals). For breeder transition, we considered all individuals who survived their first year from the same cohorts as our breeder LRS metric (1991-2014) to have complete breeding histories (*N* = 1,270 individuals). We considered an individual to have successfully established as a breeder if they were the dominant breeder on a territory during the breeding season at least once in their lifetime. Collectively, the sequence of juvenile survival, breeder transition, and breeder survival and ARS encompass all components of LRS from reaching day 11 of age to producing day 11 offspring.

### Fitness models

To determine the impact of immigrant ancestry on individual fitness, we fitted a Bayesian generalized linear mixed-effect model (GLMM) for each fitness metric with fixed effects of ancestry category, sex, and category x sex interactions. Ancestry categories were factors of eight levels in all adult models (residents, F1, F2, immigrant backcrosses, resident backcrosses, immigrants, immigrant offspring, and uncertain grandparents). The juvenile survival model included only individuals hatched on the study tract, thus excluding the immigrant and uncertain grandparent categories.

We included additional fixed effects to better describe variation in each fitness metric. Both breeder survival and ARS models included linear and quadratic terms for individual age to capture the increase in survival probability as a breeder matures and subsequent decrease at older ages due to senescence (Woolfenden and Fitzpatrick 1984). The juvenile survival model included hatch date to account for seasonal effects (Woolfenden and Fitzpatrick 1984; Barve et al. 2025). Random effects included parental identity and cohort year for LRS; parental identity, year, and individual identity for adult survival and ARS; and parental identity, year, and nest identity (to account for correlated survival among nestlings) for juvenile survival. We set the parental identity of immigrant founders with unknown parents to be unique for each individual. We fitted our LRS and ARS models using a hurdle model following (Hadfield 2014) to account for zero-inflation.

We fit all models with the MCMCglmm package (Hadfield 2024) using uninformative Gaussian priors (mean = 0, variance = 1,000) and degree-of-belief parameter (nu) of 1 for the binomial models and 0.002 for the hurdle models. We initiated our models using 1,260,000 iterations, with 60,000 burn-in and a thinning interval of 300. We increased these parameters up to a maximum of 8,200,000 iterations, 200,000 burn-in, and a thinning interval of 1000 as needed for all models to converge with effective sample sizes > 900 for all fixed effects. Our juvenile survival model was run using our initial parameter set, with the annual reproductive success model run on the maximum parameter set. We ran 3 independent chains for each model. We visually examined autocorrelation and used the “heidel.diag” and “gelman.plot” functions from the coda package (Plummer et al. 2006) to check the Heidelberger and Welch’s convergence diagnostic (Heidelberger and Welch 1983) and the Gelman-Ruben shrink factor (Brooks and Gelman 1998) for all fixed and random effects. We report model results after back-transformation to the phenotypic scale for ease of interpretation. We applied these back-transformations to the full posterior distributions for all estimates to propagate uncertainty. Summaries of all models can be found in Supplementary Materials Tables S1-5.

### Line-cross comparisons

We tested for the effects of heterosis and epistasis by calculating fitness differences between specific categories based on line-cross theory, per (Dickel et al. 2023). If variation in fitness is primarily due to additive genetic variation, individuals should exhibit the mean fitness of their parental categories (Mather and Jinks 1978; Roff and Emerson 2006). Thus, deviations in F1 fitness from the mean fitness of immigrants and residents indicate that non-additive effects such as heterosis or genetic incompatibilities influence F1 fitness. These deviations should be halved in F2 individuals because they should be homozygous for immigrant or resident ancestry at 50% of loci on average. Deviations in F2 fitness from the mean of F1 and the immigrant-resident mean fitness indicate there are additional factors beyond additive and dominance effects, namely epistasis (Dickel et al. 2023). Our full set of planned comparisons is found in Table 1. We computed the posterior distributions of fitness differences using the latent mean fitness metrics for each immigrant ancestry category output by each iteration of our Bayesian models to avoid issues with uneven sample sizes and propagate uncertainty throughout our analyses. Due to the nature of the link functions used in our models, fixed effects that function additively on the latent scale of our models will behave non-additively at the phenotypic scale (de Villemereuil et al. 2016). For this reason, we present our results and the 95% confidence intervals at the latent scale in Supplementary Materials Figures S2-8 and the directly observed fitness metrics in Figure S9.

**Table 1.** List of planned line-cross comparisons and the genetic effects detected by each comparison.

| Abbreviation | Description | Interpretation |
| --- | --- | --- |
| Rbc VS $\mu(F1, R)$ | Fitness differences between resident backcross individuals and the mean of F1 and resident individuals | Deviations are caused by dominance effects not detected in F1 individuals or epistatic effects |
| F1 VS $\mu(R, IO)$ | Fitness differences between F1 individuals and the mean of residents and immigrant offspring | Deviations are caused by dominance effects |
| F2 VS $\mu(F1 \mu(R, IO))$ | Fitness differences between F2 individuals and the mean of F1 and the mean of residents and immigrant offspring | Deviations are caused by dominance effects not detected in F1 individuals or epistatic effects |
| Ibc VS $\mu(F1, IO)$ | Fitness differences between immigrant backcross individuals and the mean of F1 and immigrant offspring individuals | Deviations are caused by dominance effects not detected in F1 individuals or epistatic effects |
| IO VS R | Fitness differences between immigrant offspring individuals and resident individuals | Deviations are caused by additive effects |
| IO VS IF | Fitness differences between immigrant offspring individuals and immigrant founder individuals | Deviations are caused by environmental effects |

### Estimation of composite genetic effects

We further investigated the genetic architecture of each fitness metric by using line cross analysis to estimate composite genetic effects (CGEs), or the cumulative effect of additive, dominant, and epistatic variation across all loci that influence the focal trait (Cockerham 1954; Hill 1982; Lynch 1991). Line cross analysis uses linear models to estimate how much a given phenotype is determined by different additive, dominance, and epistasis effects, with the number of CGEs one can estimate determined by the number of different crosses (ancestry categories) available (Cockerham 1954; Hill 1982; Blackmon and Demuth 2016). Here, we followed (Lynch 1991) and considered two-locus interactions between additive loci (additive X additive epistasis), dominant loci (dominant X dominant epistasis), and interactions between additive and dominant loci (additive X dominant epistasis):

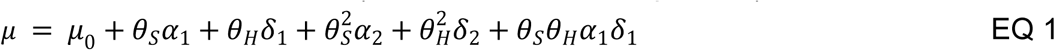

Where 𝜇 is the mean phenotype for an immigrant ancestry category

𝜇_0_ is the standard mean phenotype, in our case the phenotype of F2 individuals
𝜃_𝑆_ is the source index, a measure of degree of immigrant ancestry scaled from -1 to 1 available in Table 2
𝜃_𝐻_ is the hybridity index, a measure of degree of admixture scaled from -1 to 1 available in Table 2
𝛼_1_ is the estimate for addictive CGEs
𝛿_1_ is the estimate for dominance CGEs
𝛼_2_ is the estimate for additive X additive epistatic CGEs
𝛿_2_ is the estimate for dominance X dominance epistatic CGEs
𝛼_1_𝛿_1_ is the estimate for additive X dominance epistatic CGEs

**Table 2.** Values for the source index (𝜃_𝑆_) and hybridity index (𝜃_𝐻_) for each immigrant ancestry category used for estimating CGEs.

|  | Resident | Resident<br>Backcross | F1 | F2 | Immigrant<br>Backcross | Immigrant<br>Offspring |
| --- | --- | --- | --- | --- | --- | --- |
| $\theta_S$ | -1 | -0.5 | 0 | 0 | 0.5 | 1 |
| $\theta_H$ | -1 | 0 | 1 | 0 | 0 | -1 |

Our values for the source index and hybridity index for each immigrant ancestry category included in our analysis is located in Table 2. We used F2s as the reference population (for calculating the standard mean phenotype 𝜇_0_) (Cockerham 1954; Hill 1982) and immigrant offspring as the pure immigrant ancestry group to exclude additional environmental variation. This model assumes that all unlinked loci in the F2 generation are in Hardy-Weinberg equilibrium with respect to the parental populations. Given known inbreeding within our pedigree, we note that our models may underestimate dominance variation and both dominance X dominance and additive X dominance epistatic variation (Lynch 1991). To deal with uneven sample sizes and propagate error, we solved Equation 1 with the posterior distribution of latent mean fitness metrics for each immigrant ancestry category output by our Bayesian models to generate posterior distributions for each CGE.

### Heterozygosity

We used genomic data to assess levels of heterozygosity and pairwise relatedness between immigrants, residents, and their descendants. A previous study generated genotype data for 3,583 individuals at 15,416 single nucleotide polymorphisms (SNPs) using custom Illumina iSelect BeadChips (Chen et al. 2016). Here, we used 7,734 autosomal SNPs pruned for high linkage disequilibrium in 2,361 individuals from our study tract and time period. We used PLINK (Purcell et al. 2007) to estimate per-site heterozygosity for each individual (option -het) and to calculate pairwise relatedness between individuals (option -genome). We used the Wilcoxon signed-rank test to test for differences in pairwise relatedness between immigrants and residents and the Kruskal-Wallis rank sum test to test for overall differences in heterozygosity among ancestry groups. To directly test for the effect of heterozygosity on fitness, we fitted Bayesian GLMMs with the same fixed and random effects as before, except we replaced the immigrant ancestry category with a term for observed heterozygosity. Ungenotyped individuals (*N* = 1,653 individuals) did not contribute to the estimation of the respective fixed effects, but were retained in the models to accurately calculate the additional fixed and random effects. Summaries of all models can be found in Supplementary Materials Tables S6-10.

## Results

### Lifetime Reproductive Success

Breeder LRS was significantly greater for all females with at least 50% immigrant ancestry compared to residents (Fig 1A). Immigrant backcross females had the greatest posterior mean lifetime reproductive success (8.23 offspring), followed by F1, F2, and immigrant offspring females (Supp. Table S11; Fig 1A). For males, resident backcross individuals had the greatest LRS (8.22 offspring; Supp. Table S11; Fig 1A). Line-cross comparisons between specific groups indicated the presence of both additive and non-additive effects. Female immigrant offspring produced more offspring compared to resident females across 98% of the posterior distribution. Immigrant backcross females were less likely to produce zero offspring compared to the F1-immigrant mean (97% of the posterior distribution for this difference was negative), and both male and female F1 individuals produced more offspring compared to the immigrant-resident mean (93% of the posterior distribution was positive; Supp. Table S12). We found no difference in the lifetime reproductive success of immigrant founders and immigrant offspring, suggesting little impact of environmental effects prior to an individual arriving on the study tract (83% and 69% of the posterior distributions for the binomial and Poisson sub-models were positive; Supp. Table S12).

**Figure 1.**
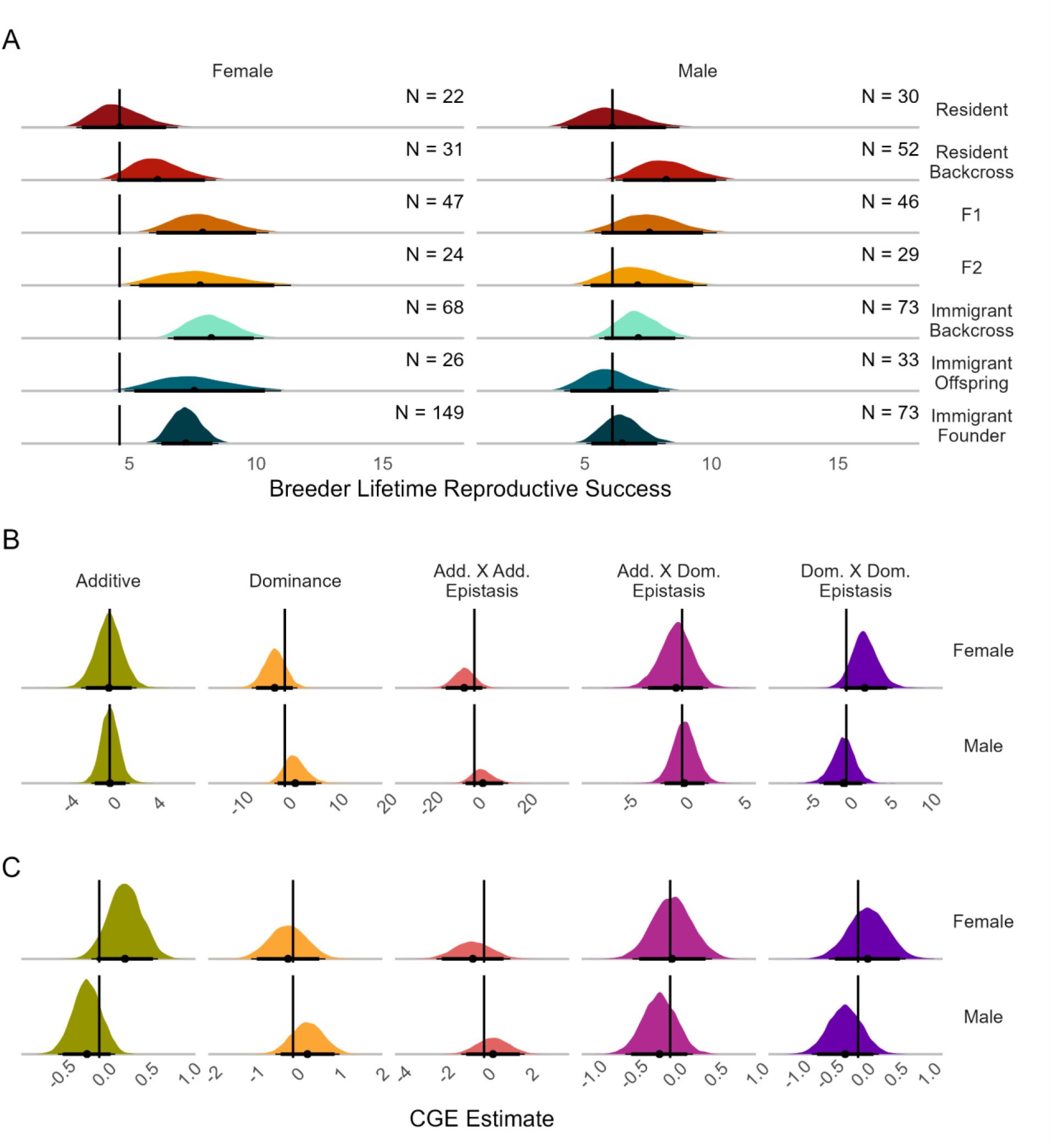
(A) Posterior distributions for breeder lifetime reproductive success at the phenotypic scale and estimates for composite genetic effects (CGEs) of the (B) binomial and (C) poisson sub-models. The posterior mean is identified with a point, with thin lines representing the 95% credible interval, and thick lines indicating the 90% credible interval. In A, the number of individuals of each sex and ancestry category (*N*) is displayed in the upper right and vertical black lines indicate the resident posterior mean. Results using the latent scale are presented in Supp. Fig S2-3. In B and C, vertical black lines indicate a CGE estimate of zero.

We then performed line-cross analysis to estimate CGEs for the probability of producing any offspring and the total number of offspring produced in males and females. We did not find any significant CGEs at the 95% or 90% credible interval for the probabilities of producing zero offspring in either sex, but found some support for dominance X dominance effects in females (85% credible interval did not overlap 0; Supp. Table S13; Fig 1B). Similarly, while all CGEs for the number of offspring produced failed to exclude zero, we found some evidence for additive effects in females (86% credible interval did not overlap 0; Supp. Table S14; Fig 1C). Collectively, our results suggest there are female-specific additive benefits of immigrant ancestry for LRS and possible heterosis in admixed individuals of both sexes driven by non-additive effects.

### Annual Breeder Fitness

To disentangle the specific components underlying the observed differences in LRS, we characterized the effects of immigrant ancestry on breeder survival and annual reproductive success. Female F1, immigrant backcross, immigrant offspring, and immigrant founders had greater annual breeder survival than residents (Supp. Table S15; Fig 2A). Annual breeder survival in males did not vary depending on immigrant ancestry. Line-cross comparisons did not find any evidence of non-additive effects (Supp. Table S16; Fig S4B). Female immigrant offspring had higher breeder survival compared to resident females (93% of the posterior distribution was positive; Supp. Table S16). Estimation of CGEs found no evidence of significant additive effects of immigrant ancestry for breeder survival in females, although female-specific additive effects had the greatest support among all breeder survival CGEs (68% credible interval did not overlap 0; Table S17; Fig 2B). Overall, our results are consistent with additive benefits of immigrant ancestry for female annual breeder survival, while immigrant ancestry had little effect on male annual breeder survival.

**Figure 2.**
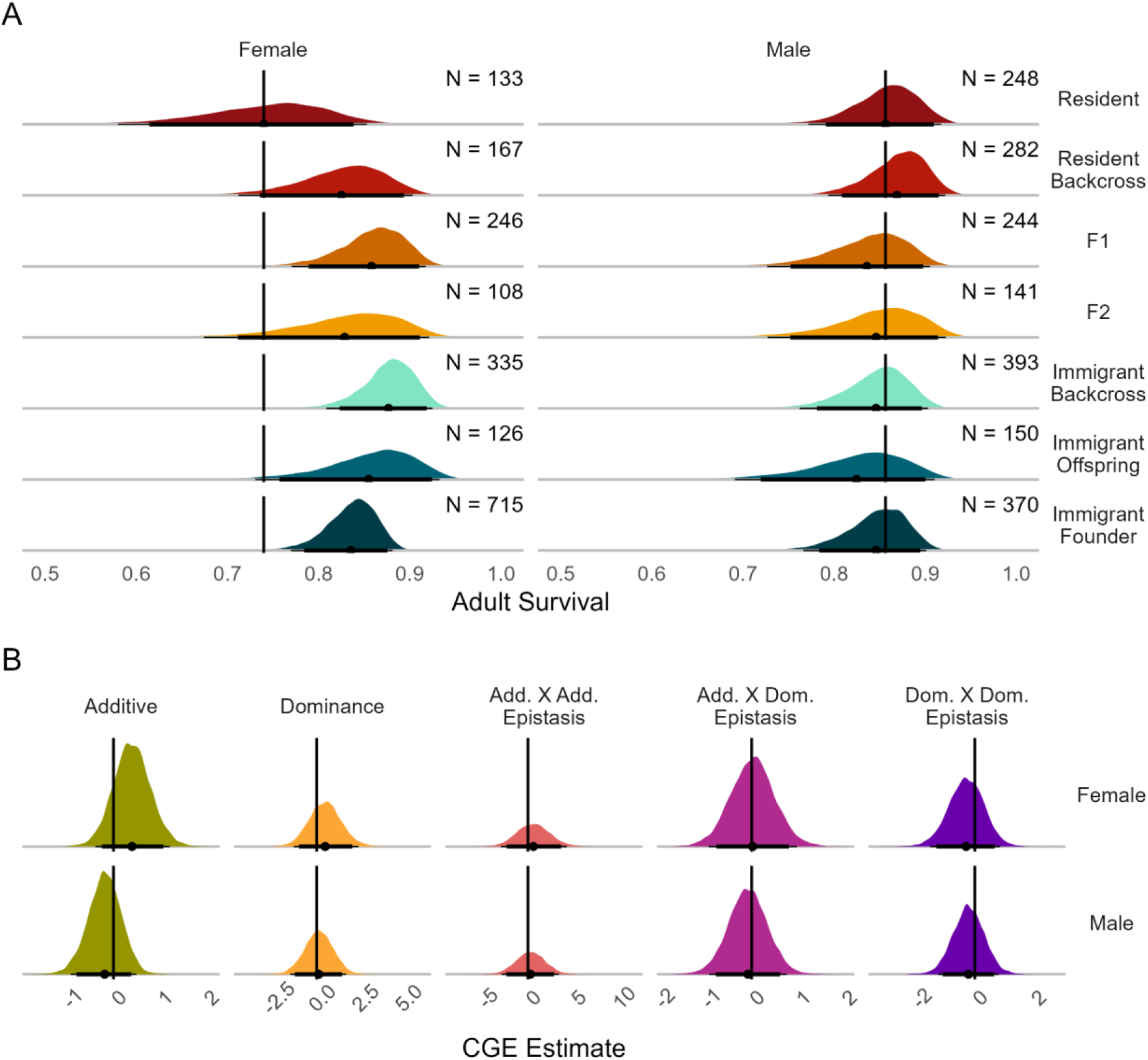
The posterior mean is identified with a point, with thin lines representing the 95% credible interval, and thick lines indicating the 90% credible interval. In A, the number of individuals of each sex and ancestry category (*N*) is displayed in the upper right and vertical black lines indicate the resident posterior mean. Results using the latent scale are presented in Supp. Fig S4. In B, vertical black lines indicate a CGE estimate of zero.

For ARS, we found no significant differences among ancestry groups, largely due to the width of the posterior distribution for each ancestry group (Supp. Table S18; Fig 3A). However, specific comparisons of ancestry groups showed that F1 males were less likely to produce zero offspring relative to expectations (93% of the posterior difference between F1s and the mean of residents and immigrant offspring was negative; Supp. Table S19; Fig S5B), providing some support for dominance effects. Immigrant offspring females were less likely to produce zero offspring relative to immigrant founders (93% of the posterior difference was negative; Supp. Table S19; Fig S5B), indicating possible environmental or social costs to immigrant founders. Estimation of CGEs provided no significant results, although the CGEs with the greatest support were additive effects for both males (83% credible interval did not overlap 0) and females (74% credible interval did not overlap 0) for the likelihood of producing no offspring, however the signs of these effects were opposite, indicating benefits in females and costs in males (Supp. Table S20-21; Fig 3B). Together, our results suggest that different fitness components contribute to variation in LRS in different sexes, with some evidence for additive benefits of immigrant ancestry for breeder survival and ARS in females and possible heterosis for ARS in males.

**Figure 3.**
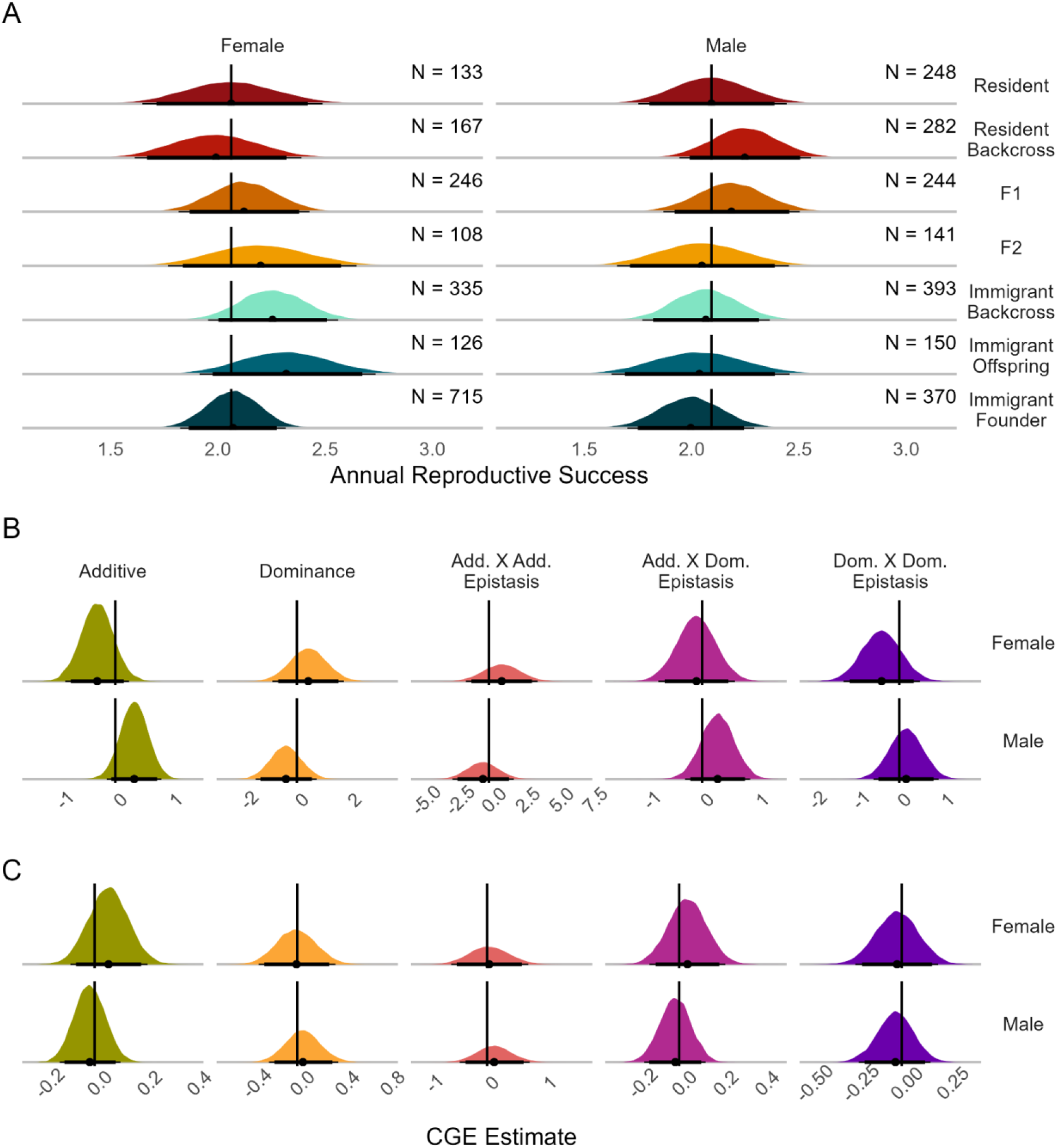
(A) Posterior distributions for breeder annual reproductive success at the phenotypic scale and model average estimates for composite genetic effects (CGEs) of the binomial (B) and poisson (C) sub-models. The posterior mean is identified with a point, with thin lines representing the 95% credible interval, and thick lines indicating the 90% credible interval. In A, the number of individuals of each sex and ancestry category (*N*) is displayed in the upper right and vertical black lines indicate the resident posterior mean. Results at the latent scale presented in Supp. Fig S5-6. In B and C, vertical black lines indicate a CGE estimate of zero.

### Juvenile Survival

Contrary to our results for breeder fitness measures, we found evidence that immigrant ancestry reduced juvenile survival compared to residents. Female resident backcross, F1, F2, and immigrant offspring individuals had lower juvenile survival than female residents (Supp. Table S22; Fig 4A). Male F1 and immigrant offspring tended to have lower juvenile survival probabilities than residents (91% and 86% of posterior distribution less than the resident mean, respectively; Fig 4A). When contrasting juvenile survival estimates to the mid-parental means, we found F2 males (91% of the posterior distribution is positive) and immigrant backcross juveniles of both sexes (94% and 91% of the posterior difference is positive for males and females, respectively) had significantly greater survival than expected (Supp. Table S23; Fig S6B). Female immigrant offspring had significantly lower juvenile survival compared to resident females (Supp. Table S23; S6B). CGE estimation found positive additive X dominant epistasis in female juvenile survival (92% credible interval did not overlap 0; Table S24; Fig 4B). These results suggest there are non-additive effects of immigrant ancestry on juvenile survival that act in opposite directions in different sexes, producing outbreeding depression in females and heterosis in males.

**Figure 4.**
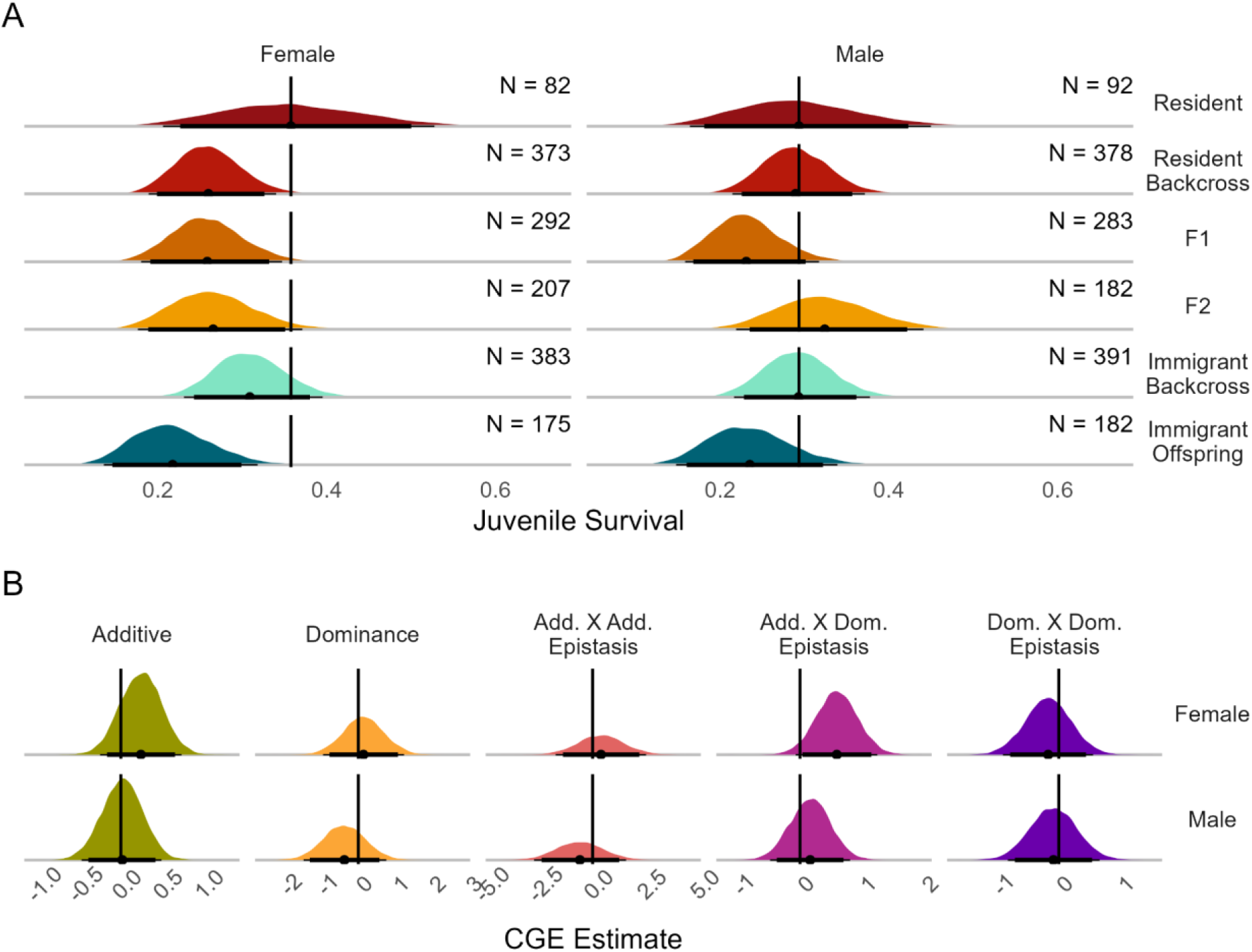
(A) Posterior distributions for juvenile survival at the phenotypic scale and (B) model average estimates for composite genetic effects (CGEs). The posterior mean is identified with a point, with thin lines representing the 95% credible interval, and thick lines indicating the 90% credible interval. In A, the number of individuals of each sex and ancestry category (*N*) is displayed in the upper right and vertical black lines indicate the resident posterior mean. Results at the latent scale presented in Supp. Fig S7. In B, vertical black lines indicate a CGE estimate of zero.

### Breeder Transition

We found that immigrant backcross females were more likely than resident females to become a breeder during their lifetime (Supp. Table. S25; Fig. 5A). We also see a dramatic increase in probability of becoming breeders for immigrant founders of both sexes, which likely reflects a limitation of our data (immigrants who do not stay in our study population for multiple months are less likely to be detected; Supp. Table S26). Comparisons of specific ancestry groups found some evidence of non-additive effects, with resident backcross males having a greater than expected likelihood of becoming breeders (90% of the posterior distribution was greater than the F1-resident mean; Supp. Table S26). These results were supported by the line-cross analysis, which found the strongest support for negative dominance X dominance epistatic effects in males (70% credible interval did not overlap 0; Table S27; Fig. 5B), although none of the estimated CGEs significantly diverged from zero. Thus, for the probability of transitioning to breeder status given survival to their first year, there may be non-additive benefits of admixture in males.

**Figure 5.**
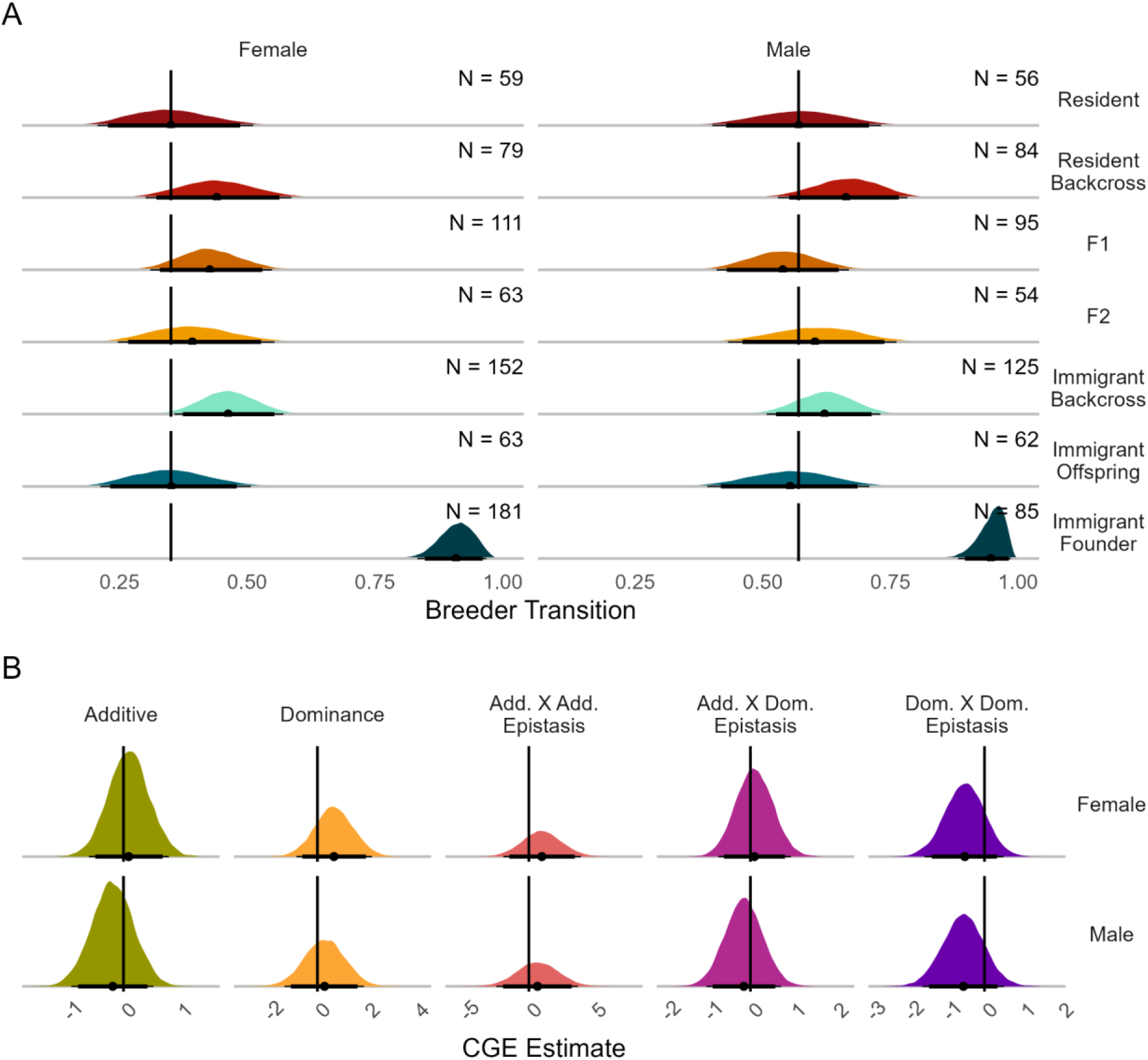
(A) Posterior distributions for lifetime breeder transition rate at the phenotypic scale and (B) model average estimates for composite genetic effects (CGEs). The posterior mean is identified with a point, with thin lines representing the 95% credible interval, and thick lines indicating the 90% credible interval. In A, the number of individuals of each sex and ancestry category (*N*) is displayed in the upper right and vertical black lines indicate the resident posterior mean. Results at the latent scale presented in Supp. Fig S8. In B, vertical black lines indicate a CGE estimate of zero.

### Heterozygosity

We used genomic data to characterize differences in relatedness and heterozygosity among immigrant ancestry categories. Pairwise relatedness among immigrants was lower than relatedness among residents (Wilcoxon test: W = 1.5x10^10, *p* < 0.0001), likely because immigrants come from multiple source populations. Levels of heterozygosity were different based on ancestry category (Kruskal-Wallis test: 𝜒^2^ = 41.818, *p* < 0.0001), but only immigrant backcross individuals showing significantly different heterozygosity relative to residents after correcting for multiple tests (resident mean = 0.347, immigrant backcross mean = 0.349; Wilcoxon rank-sum test: *W* = 97213, *p* = 0.0027; Fig S11). Models of heterozygosity effects on fitness found sex-specific effects of heterozygosity on juvenile survival and likelihood to become a breeder (Supp. Fig. S10). More heterozygous females were more likely to survive their first year (Supp. Fig. S10A), and more heterozygous male immigrant founder individuals were more likely to become breeders during their lifetime (Supp. Fig. S10F). Overall, these results indicate that the costs of being inbred may only be strong enough to detect during particular life stages for specific individuals in the population.

## Discussion

We investigated the multigenerational fitness outcomes of gene flow in a population with historically high immigration by comparing the fitness of immigrants and their descendants using line-cross theory. We found benefits of immigrant ancestry that persisted over multiple generations, resulting in peak fitness of immigrant backcross individuals. We then explicitly estimated the cumulative effects of additive, dominant, and epistatic loci, and found evidence of non-additive effects driving heterosis in adults while contributing to outbreeding depression in juveniles. These patterns were likely driven by dominance X dominance and additive X dominance epistasis, respectively, indicating that novel interactions between loci likely play a role in the fitness of immigrant descendants. Understanding the complex genetic effects of dispersal is vital for predicting the outcomes of gene flow as habitat fragmentation increasingly isolates populations.

The increased lifetime reproductive success of all immigrant ancestry categories relative to residents in females can likely be attributed to a combination of additive benefits and positive heterosis. Strong positive heterosis is expected in highly inbred populations where outbreeding rescues the population from the effects of inbreeding depression (Tallmon et al. 2004; Frankham 2015). While inbreeding depression is observed in our focal population (Chen et al. 2016), the high amount of gene flow (averaging 10 immigrants per year) far exceeds the expected amount required to maintain minimal divergence (1-10 immigrants per generation) (Mills and Allendorf 1996), making it surprising to discover a measurable degree of heterosis. Further, we found that the benefits of heterosis existed for F2 and immigrant backcross individuals in addition to F1s. While heterosis typically leads to the greatest fitness gains within F1 individuals (Whitlock et al. 2000; Marr et al. 2002; Charlesworth and Willis 2009), heterosis driven by epistatic interactions instead of dominance alone will appear in later generations, where novel interactions across loci can emerge (Lynch 1991; Costa e Silva et al. 2012; Le Rouzic et al. 2024). These interactions include synergistic masking of deleterious variation due to dominance X dominance epistatic effects, which we detected for female LRS. Recurrent outbreeding can also maintain heterozygosity over multiple generations (Whitlock et al. 2000; Johnson et al. 2011; Fitzpatrick et al. 2020), increasing the rate of introgression because the most outbred lineages reproduce faster than lineages with more resident ancestry (Ingvarsson and Whitlock 2000). The high levels of immigration into ABS maintain ample opportunities for recurrent outbreeding, matching many successful cases of genetic rescue through repeated translocations (Frankham 2015; Poirier et al. 2019; Onorato et al. 2024). Immigrants into ABS also likely originate from different sub-populations, potentially increasing the level of admixture of the individuals with high immigrant ancestry, leading to nearly equivalent heterozygosity among immigrant backcross individuals as F1s. These results reinforce the viability of increasing gene flow to create prolonged fitness gains in natural populations (Frankham 2015), but also emphasize that resident lineages may remain rare despite overall population stability. Our findings highlight the importance of monitoring the fitness outcomes of immigrants beyond a single generation.

We found that immigrant ancestry had a greater effect on fitness in females than in males. Sex-specific fitness benefits, or different fitness costs of dispersal are theoretically expected (Greenwood 1980; Li and Kokko 2019) and have been observed in many species (Gienapp and Merilä 2011; Green and Hatchwell 2018; Barbraud and Delord 2021). These sex-specific differences are theorized to exist due to different degrees of intrasexual competition for resources, where the sex that is more likely to gain resources via philopatry suffers greater costs of dispersal (Greenwood 1980). In the Florida scrub-jay, males are far more likely than females to inherit natal territory (Woolfenden and Fitzpatrick 1978), and individuals who inherit natal territory have greater reproductive success than all other forms of territory acquisition (Woolfenden and Fitzpatrick 1984; Fitzpatrick et al. 1999; Shah et al. 2024). While these differing costs can explain why immigrant founders outperformed residents, they cannot explain the benefits of immigrant ancestry persisting over multiple generations. One possibility is the heritability of dispersal-related traits, such as explorative behavior (Korsten et al. 2013), in immigrant lineages providing benefits to the more dispersive females and having neutral or negative effects in males. Heritable traits related to dispersal have been observed in several bird species (Hansson et al. 2003; Pasinelli et al. 2004; Korsten et al. 2013). Similarly, the recruitment of offspring who dispersed to other sub-populations contributed to the fitness of dispersing individuals in a metapopulation of house sparrow (*Passer domesticus*; Saatoglu et al. 2025), and dispersal has been shown to be heritable in this same metapopulation (Saatoglu et al. 2024), linking heritable dispersal with increased fitness among immigrants. Alternatively, the greater rate of female immigration relative to males could lead to greater additive genetic variation, allowing for the signal of positive additive effects of immigrant ancestry in females. In our population of Florida Scrub-Jays, we found significant additive genetic variation for LRS in females but not males (Cosgrove et al. 2025). The multigenerational female-specific benefits of immigrant ancestry thus could result from the introduction of beneficial alleles to the focal population from different source populations of immigrants.

Unlike adult fitness measures, we observed apparent outbreeding depression in female juvenile survival, indicating a cost of immigrant ancestry driven by epistatic effects. Florida scrub-jays rarely disperse as juveniles, thus it is unlikely that this mortality represents increased dispersal among immigrant descendants (Woolfenden and Fitzpatrick 1984). Instead, the apparent outbreeding depression could be caused by differences in group size. Florida scrub-jays live in family groups with potentially several non-breeding adult helpers, who are associated with increased juvenile survival in Florida scrub-jays (Woolfenden 1975; Woolfenden and Fitzpatrick 1984; Fitzpatrick and Bowman 2016). Nests with immigrant offspring were the least likely to have helpers present (22% of nests had helpers, compared to 33% of resident nests). Nests with immigrant backcross individuals were the most likely to have helpers (37%), and there was an overall significant effect of immigrant ancestry category on the likelihood of a nest having helpers (𝜒^2^ = 31.866, *p* < 0.001). Helpers contribute the most to juvenile survival, as compared to juvenile production and adult survival (Summers et al. 2025), possibly explaining why costs of immigrant ancestry were only apparent in juvenile survival. Yet, the difference in helper presence cannot explain the persistent lower fitness of resident backcross, F1, and F2 offspring relative to residents in females. It remains possible that there are negative effects of immigrant ancestry that are expressed in female juveniles, with the costs compensated in immigrant backcross individuals by either the high presence of helpers or the high levels of heterozygosity within this group, which also was associated with greater survival of female juveniles.

High heterozygosity of immigrant backcross individuals is consistent with our hypothesis that immigrants originate from multiple source populations. These social effects and the unexpectedly high heterozygosity of immigrant backcross individuals likely interfere with the ability of our line cross analysis to attribute differences in juvenile survival to specific CGEs.

Traditional line cross theory uses inbred parental lines to create uniform generations and control for factors such as environmental and parental effects (Miller et al. 1963; Hill 1982). Using a natural population introduces greater genetic variation that is unaccounted for in our line cross analysis, lowering our power to detect significant differences between groups, and grouping together reciprocal crosses to ensure sufficient sample sizes made it impossible to distinguish between parental effects of particular crosses. We used Bayesian models to produce estimates for generation means instead of direct measures to account for variation among cohorts and siblings, as well as variation introduced by different sample sizes for immigrant ancestry categories. We note that the values of CGEs calculated from our line cross analysis are dependent on the initial allele frequencies and non-equilibrium genotype frequencies in our sample (Mather and Jinks 1978; Lynch 1991). Thus, future analyses of the response of our focal population to selection need to account for allele frequency differences between our resident and immigrant categories and factors such as inbreeding that impact genotype frequencies. The addition of whole genome sequencing data for our study population and surrounding populations could potentially differentiate immigrant source populations and further tease apart the role of environmental factors and different genetic effects in the fitness of immigrant lineages. Theory predicts that differences in immigrant source populations can have far-reaching impacts on the fitness effects of gene flow due to differences in genetic load (Frankham et al. 2011; Mathur and DeWoody 2021; Kyriazis et al. 2023; Goedert et al. 2025).

Understanding how genetic load functions in populations that previously experienced high immigration, and how the effects of gene flow change over time during isolation, would greatly benefit efforts to predict population responses to fragmentation and to determine how and when to prioritize populations for conservation.

## Conclusions

We found persistent increased fitness among individuals with greater immigrant ancestry in a population with relatively high rates of immigration, showing that heterosis can lead to fitness benefits for multiple generations even outside of highly inbred populations. These benefits were not universal across fitness metrics, which demonstrates the importance of considering the multiple components that comprise individual fitness and the role of environmental and social factors in influencing rates of introgression. The fitness benefits of heterosis in our system were also not driven by dominance effects, as we expected, but by epistatic effects. Epistatic variation can be converted to additive variation when populations undergo bottlenecks or experience inbreeding (Lynch 1991; Cheverud and Routman 1996; Burch et al. 2024), thus the epistatic variation we observed may act as a reservoir for future additive variation. In our study population of Florida scrub-jays, immigrant backcross individuals were both extremely common and had high fitness, possibly accelerating the rate of introgression (Ingvarsson and Whitlock 2000) and leading to high genetic contributions of immigrants (Chen et al. 2019). This rapid rate of introgression and resulting increase in fitness suggests that efforts to increase connectivity may be highly successful. Research into the multigenerational fitness effects of immigration within the Florida scrub-jays at ABS sheds light on the factors that control effective gene flow more broadly, with important implications for conserving the many populations that currently face increasing habitat fragmentation due to anthropogenic change.

## Data and Code Availability Statement

All data and code used in this study can be found at https://github.com/jtsummers53/ImmigrantAncestryFitness

## Author Contributions

Jeremy Summers: Conceptualization, Methodology, Software, Resources, Formal analysis, Investigation, Data Curation, Writing - Review and Editing.

Elissa J. Cosgrove: Investigation and Data Curation. Tori Bakley: Investigation and Data Curation.

Sahas Barve: Investigation and Data Curation. Reed Bowman: Investigation and Data Curation.

John W. Fitzpatrick: Investigation and Data Curation.

Nancy Chen: Conceptualization, Methodology, Investigation, Resources, Data Curation, Writing - Review and Editing, Supervision, Project Administration, and Funding Acquisition.

## Acknowledgements

We would like to thank all the interns and staff at Archbold Biological Station who collected demographic data on the Florida Scrub-Jay over the past half-century. We thank the members of the Chen Lab and the Avian Ecology Program at Archbold Biological Station for their valuable feedback.

