## Supplementary Information for "Sustained multigenerational fitness benefits of natural immigration"

### Supplementary Figures

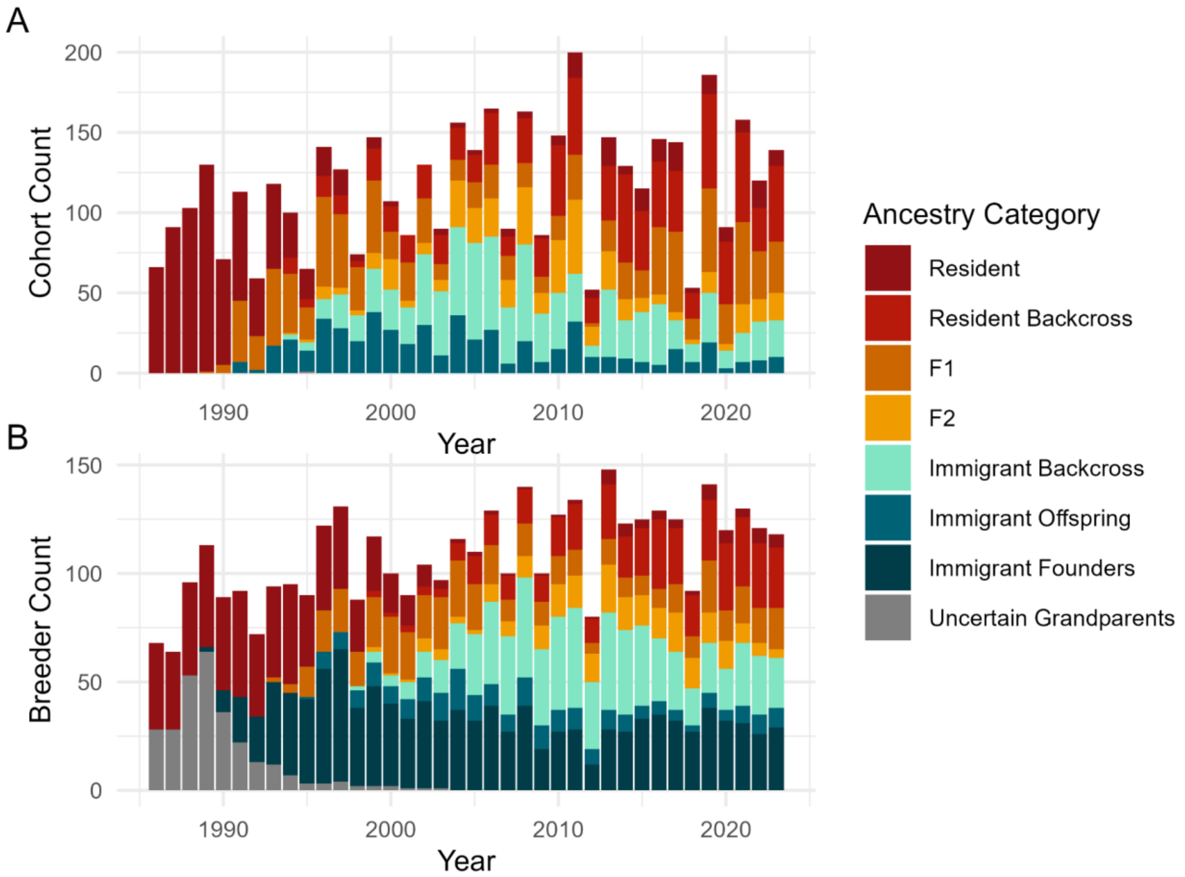


**Figure S1.** Distribution of immigrant ancestry categories among (A) the locally hatched cohort and (B) active breeders each year. Individuals are categorized based on the number of known immigrant grandparents they have: 0 for Residents, 1 for Resident Backcross, 2 for F1 and F2 (F2 individuals have two F1 parents), 3 for Immigrant Backcross, and 4 for Immigrant Offspring and Founders (Immigrant Offspring were hatched on the study tract). Individuals with uncertain grandparents could not be assigned to an immigrant ancestry category.

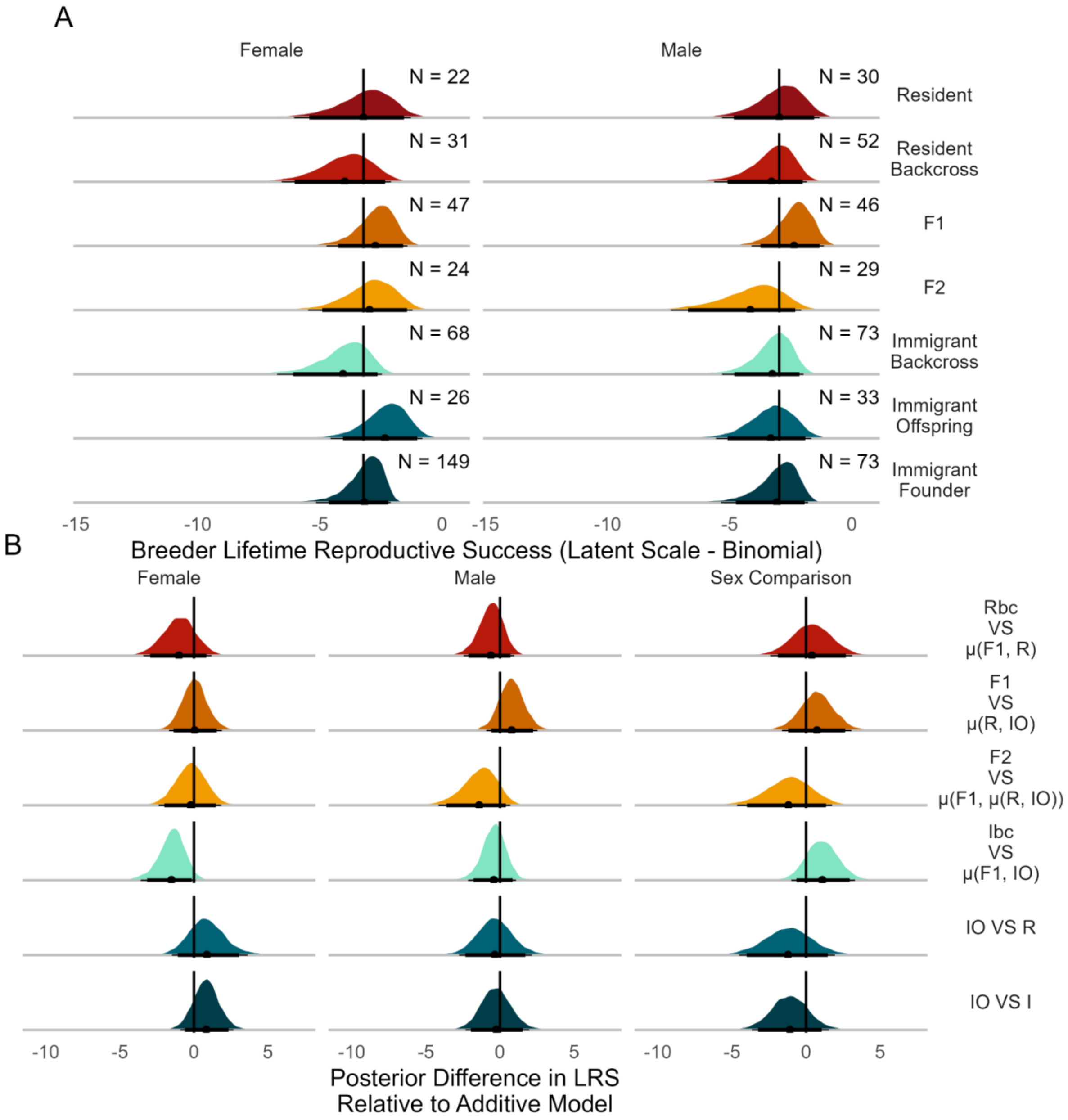


**Figure S2.** (A) Posterior distributions for breeder lifetime reproductive success, binomial sub-model for likelihood of having zero offspring, at the latent scale and (B) comparisons between ancestry categories and their mid-parental value. The posterior mean is identified with a black point, with thin lines representing the 95% credible interval, and thick lines indicating the 90% credible interval. In A, the number of individuals of each sex and ancestry category (*N*) is displayed in the upper right. In B, vertical black lines indicate zero deviations from the mid-parental value, and the right column shows fitness differences between females and males, with positive values indicating greater values in females. See Table 1 for explanations of the line-cross comparisons.

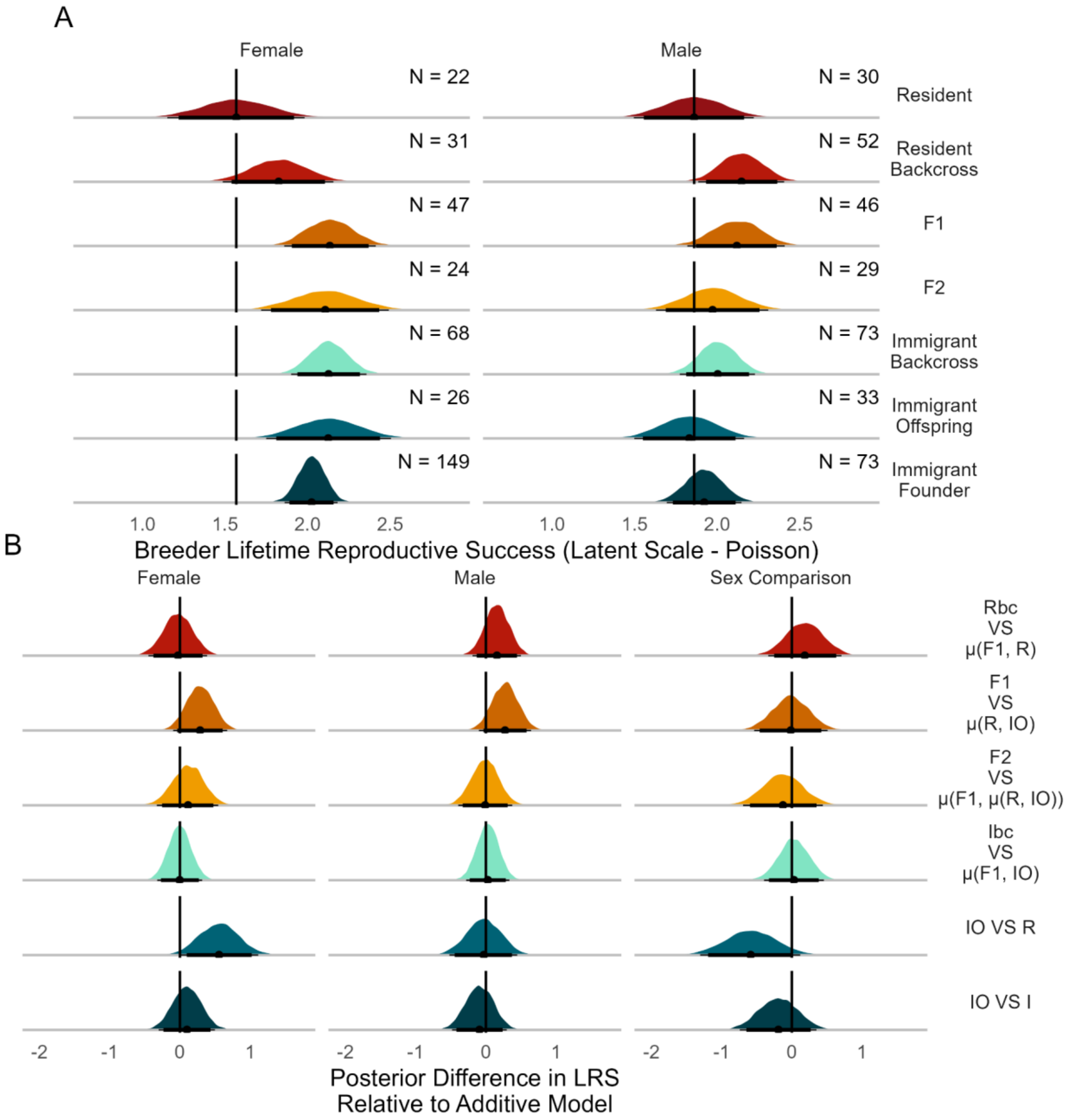


**Figure S3.** (A) Posterior distributions for breeder lifetime reproductive success, poisson sub-model for number of offspring, at the latent scale and (B) comparisons between ancestry categories and their mid-parental value. The posterior mean is identified with a black point, with thin lines representing the 95% credible interval, and thick lines indicating the 90% credible interval. In A, the number of individuals of each sex and ancestry category (*N*) is displayed in the upper right. In B, vertical black lines indicate zero deviations from the mid-parental value, and the right column shows fitness differences between females and males, with positive values indicating greater values in females. See Table 1 for explanations of the line-cross comparisons.

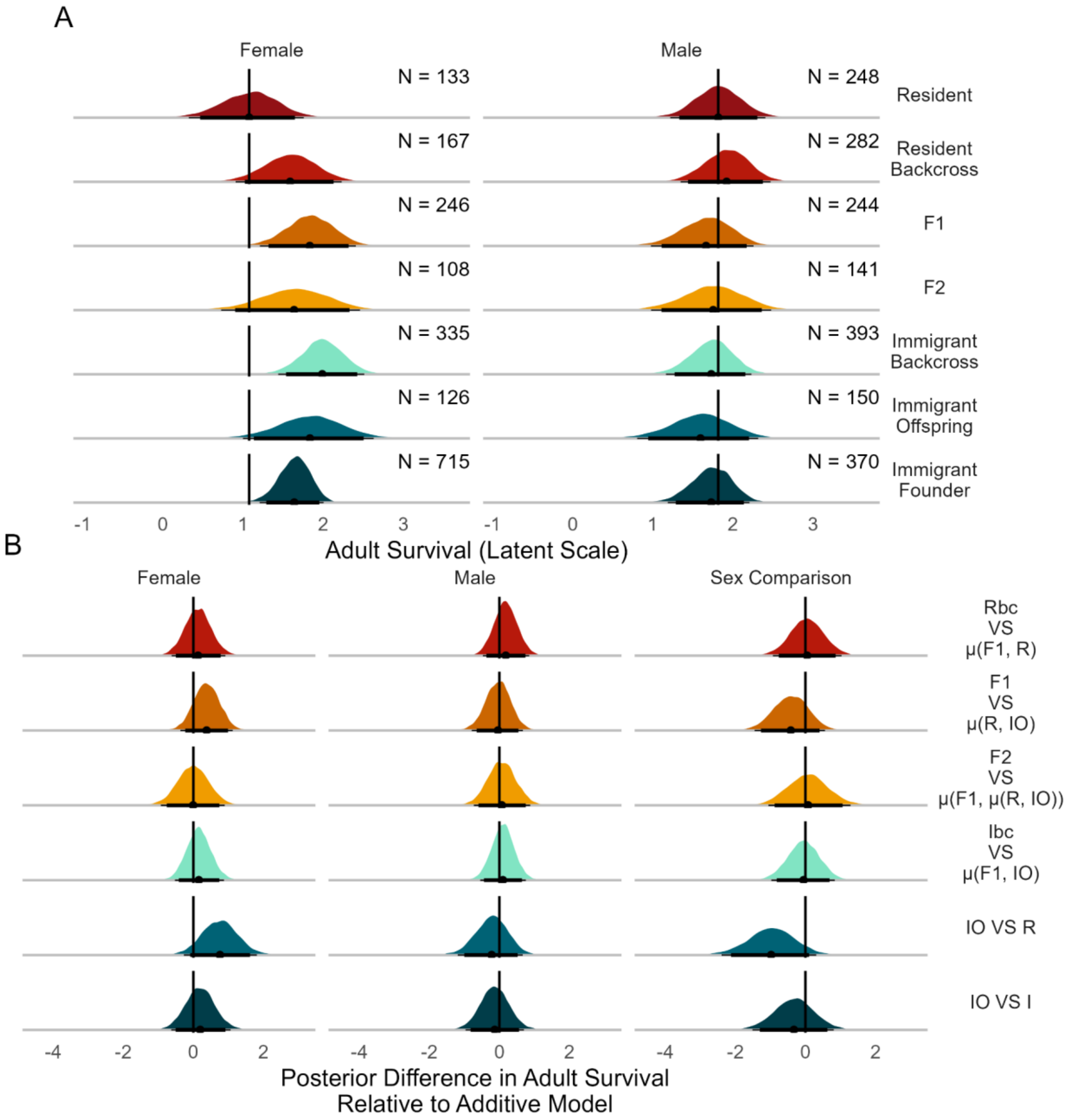


**Figure S4.** (A) Posterior distributions for breeder survival at the latent scale and (B) comparisons between ancestry categories and their mid-parental value. The posterior mean is identified with a black point, with thin lines representing the 95% credible interval, and thick lines indicating the 90% credible interval. In A, the number of individuals of each sex and ancestry category (*N*) is displayed in the upper right. In B, vertical black lines indicate zero deviations from the mid-parental value, and the right column shows fitness differences between females and males, with positive values indicating greater values in females. See Table 1 for explanations of the line-cross comparisons.

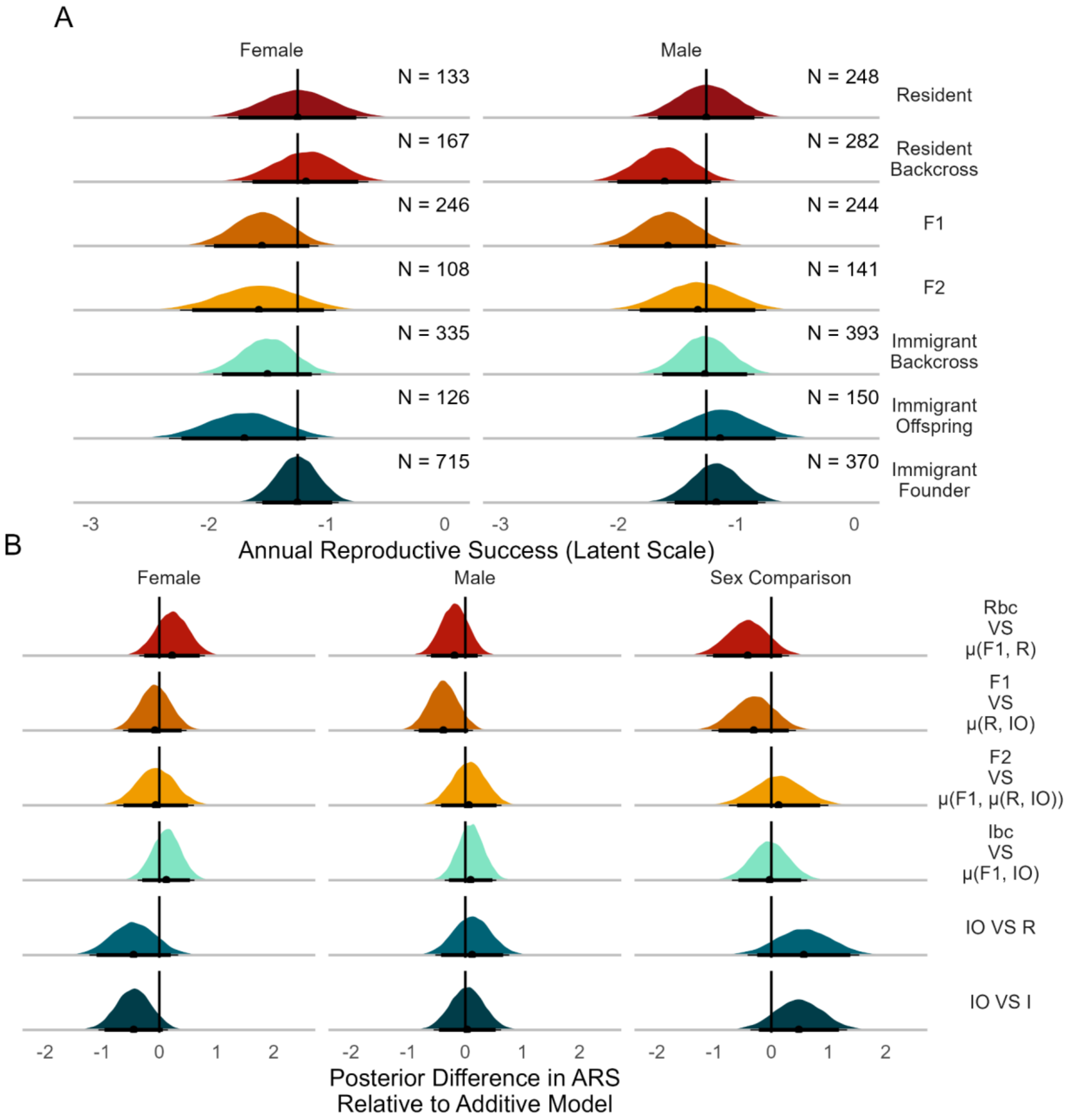


**Figure S5.** (A) Posterior distributions for annual lifetime reproductive success, binomial sub-model for likelihood of having zero offspring, at the latent scale and (B) comparisons between ancestry categories and their mid-parental value. The posterior mean is identified with a black point, with thin lines representing the 95% credible interval, and thick lines indicating the 90% credible interval. In A, the number of individuals of each sex and ancestry category (*N*) is displayed in the upper right. In B, vertical black lines indicate zero deviations from the mid-parental value, and the right column shows fitness differences between females and males, with positive values indicating greater values in females. See Table 1 for explanations of the line-cross comparisons.

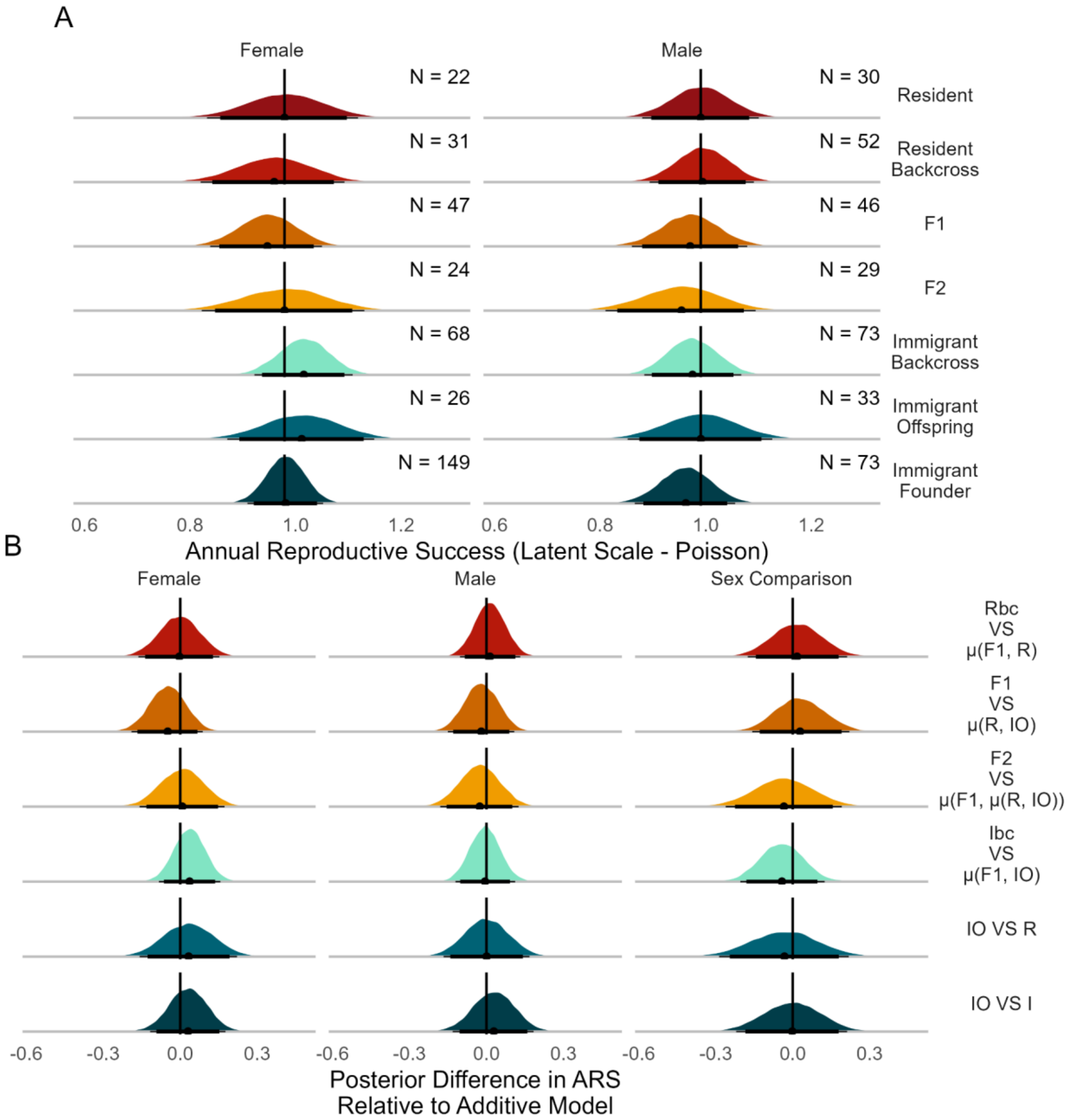


**Figure S6.** (A) Posterior distributions for annual lifetime reproductive success, poisson sub-model for number of offspring, at the latent scale and (B) comparisons between ancestry categories and their mid-parental value. The posterior mean is identified with a black point, with thin lines representing the 95% credible interval, and thick lines indicating the 90% credible interval. In A, the number of individuals of each sex and ancestry category (*N*) is displayed in the upper right. In B, vertical black lines indicate zero deviations from the mid-parental value, and the right column shows  fitness differences between females and males, with positive values indicating greater values in females. See Table 1 for explanations of the line-cross comparisons.

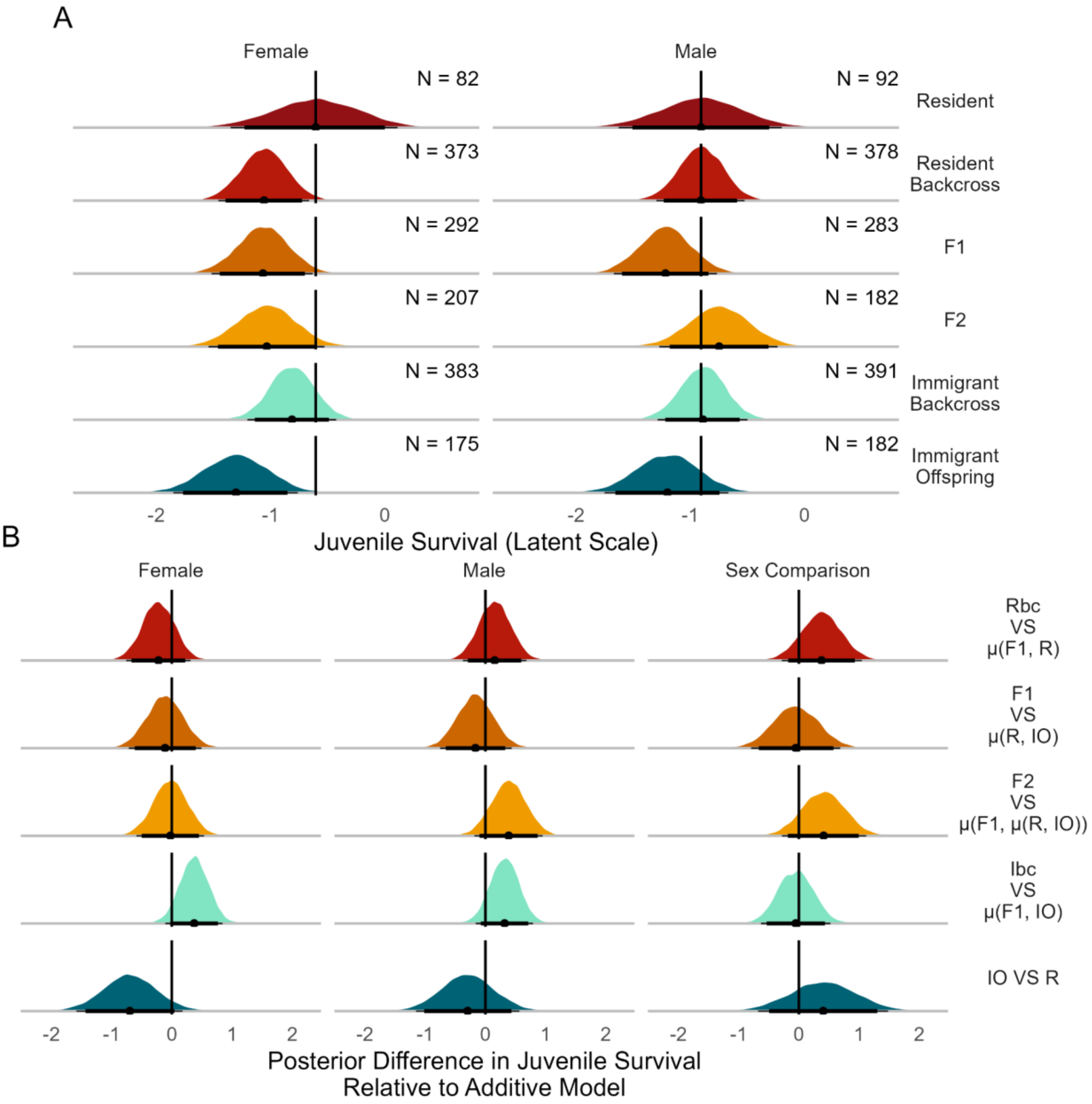


**Figure S7.** (A) Posterior distributions for juvenile survival at the latent scale and (B) comparisons between ancestry categories and their mid-parental value. The posterior mean is identified with a black point, with thin lines representing the 95% credible interval, and thick lines indicating the 90% credible interval. In A, the number of individuals of each sex and ancestry category (*N*) is displayed in the upper right. In B, vertical black lines indicate zero deviations from the mid-parental value, and the right column shows  fitness differences between females and males, with positive values indicating greater values in females. See Table 1 for explanations of the line-cross comparisons.

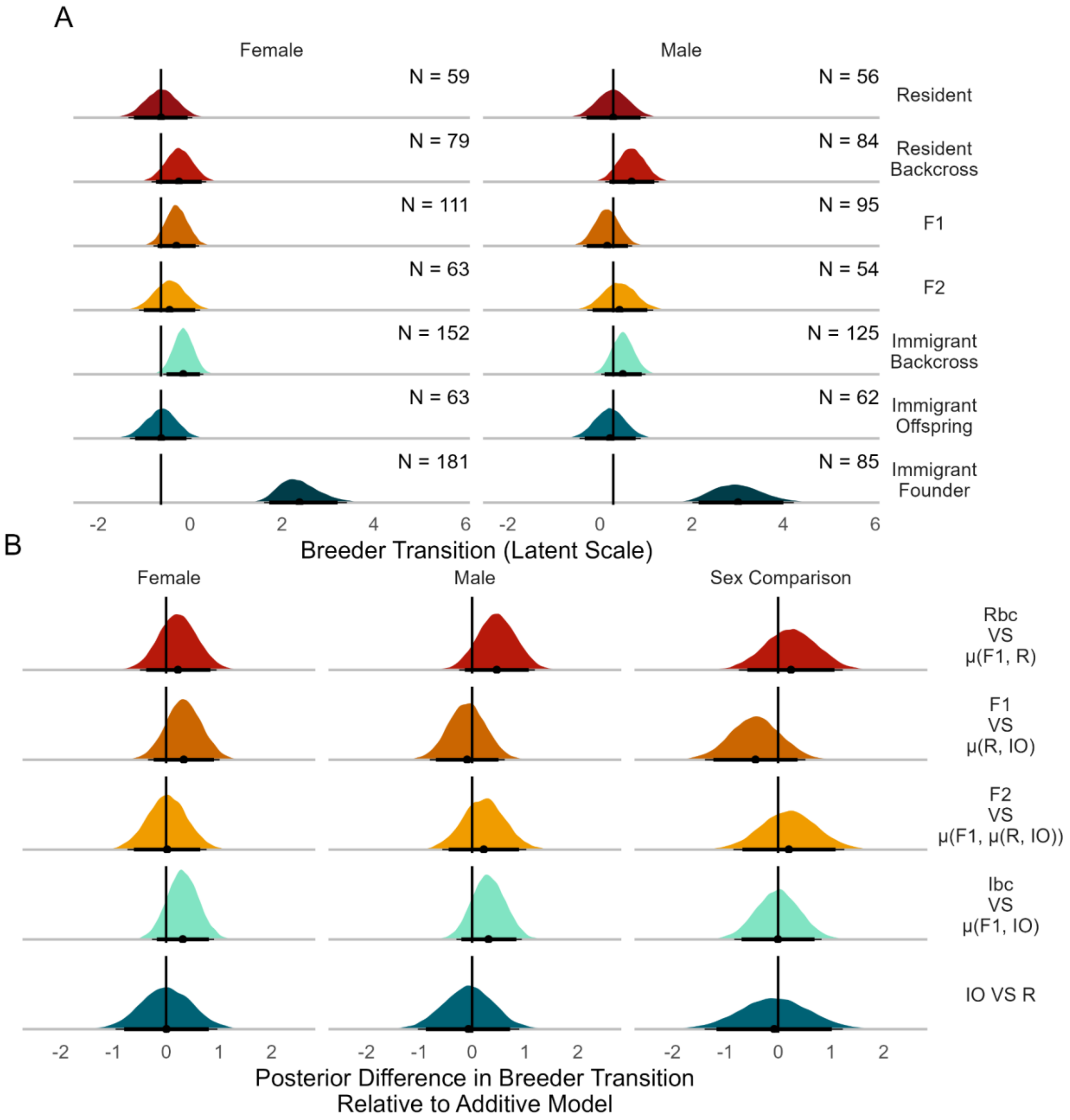


**Figure S8.** (A) Posterior distributions for likelihood of becoming a breeder at the latent scale and (B) comparisons between ancestry categories and their mid-parental value. The posterior mean is identified with a black point, with thin lines representing the 95% credible interval, and thick lines indicating the 90% credible interval. In A, the number of individuals of each sex and ancestry category (*N*) is displayed in the upper right. In B, vertical black lines indicate zero deviations from the mid-parental value, and the right column shows fitness differences between females and males, with positive values indicating greater values in females. See Table 1 for explanations of the line-cross comparisons.

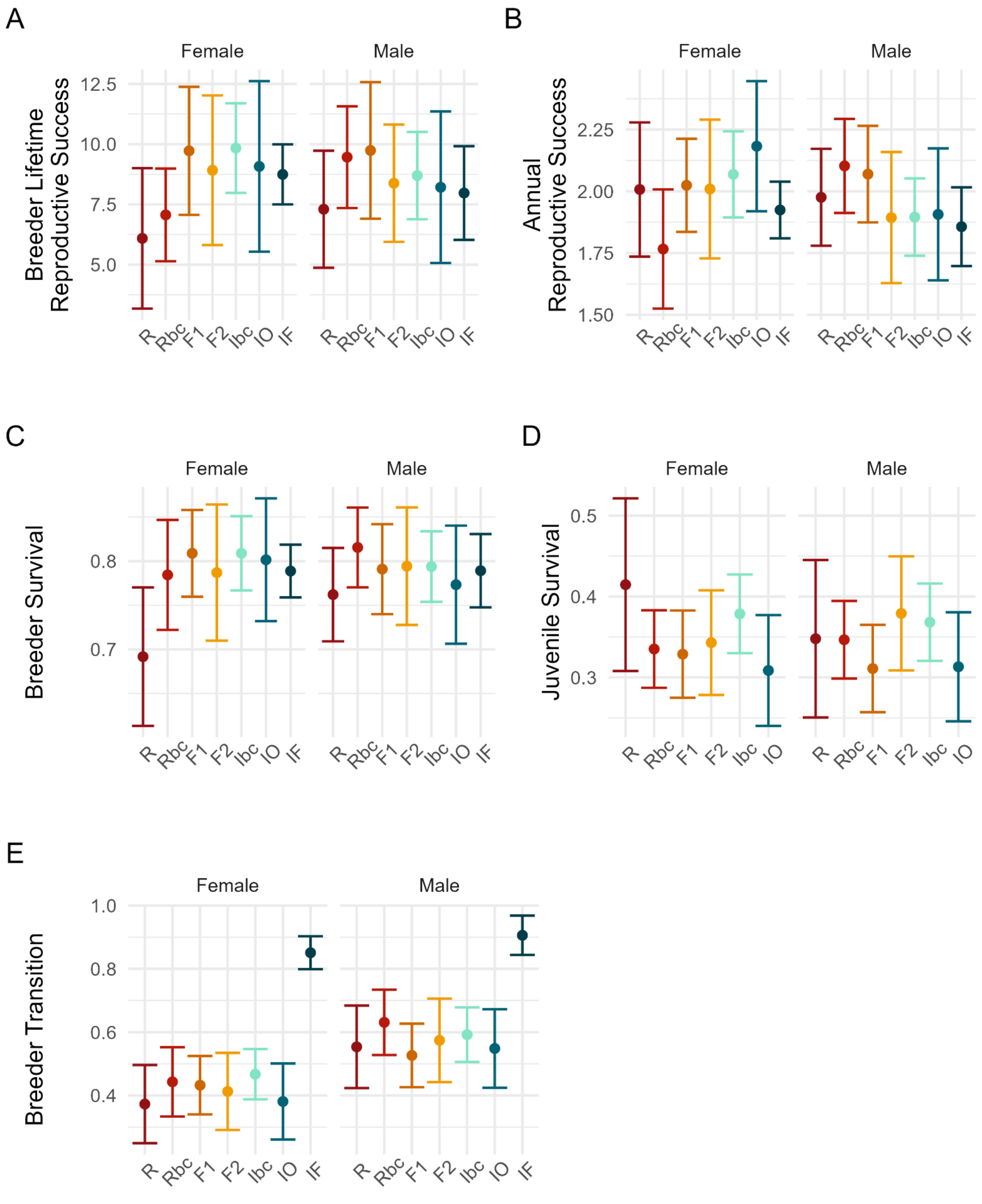


**Figure S9.** Observed means for (A) breeder LRS, (B) ARS, (C) breeder survival, and (D) juvenile survival for females and males of each immigrant ancestry category. The 95% confidence interval is denoted with errorbars. Abbreviations for ancestry categories are: R = resident, Rbc = resident backcross, F1, F2, Ibc = immigrant backcross, IO = immigrant offspring, and IF = immigrant founder.

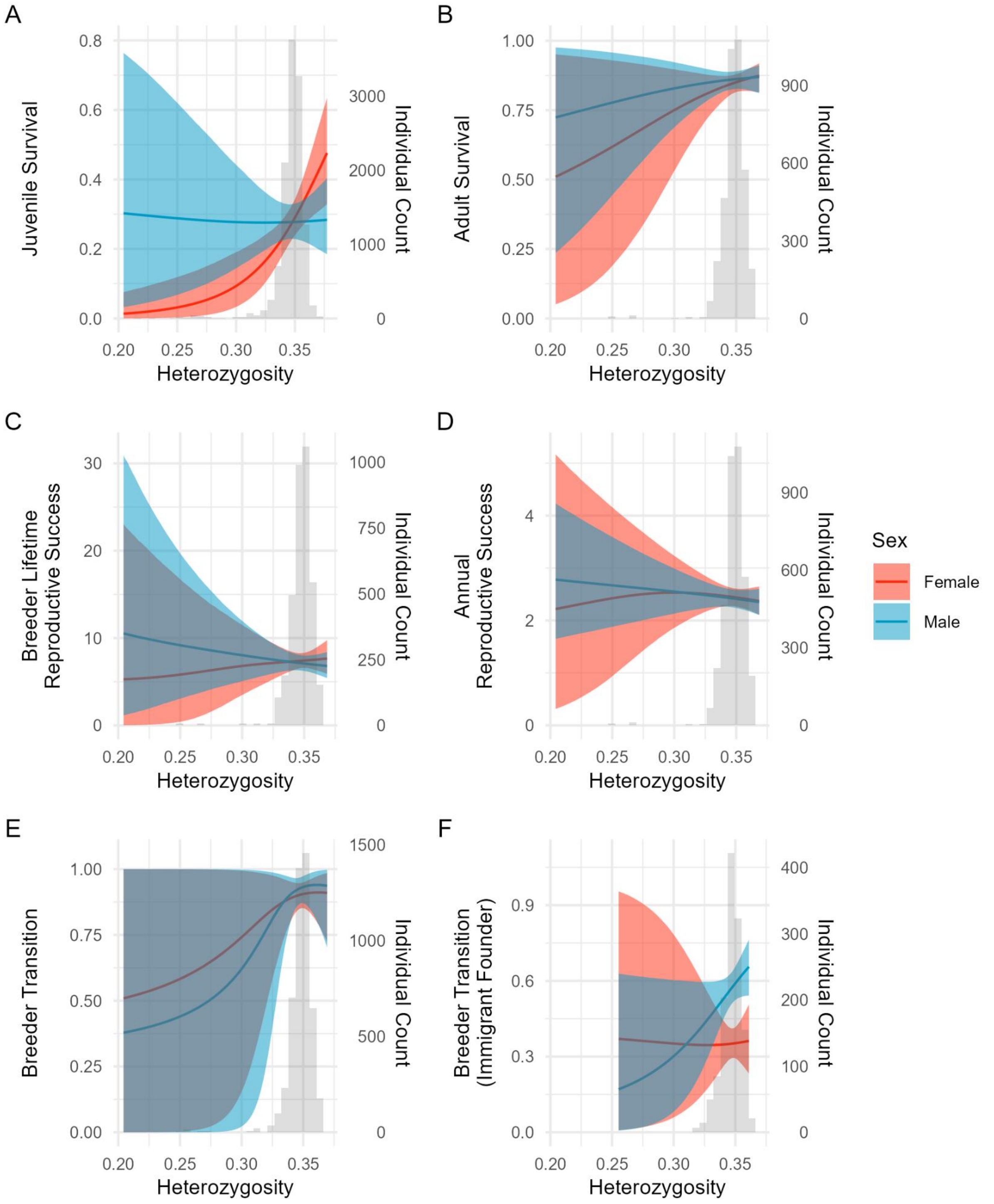


**Figure S10.** Posterior predictions for the effect of heterozygosity on different fitness metrics for females (red) and males (blue). Solid lines indicate the posterior mean, with shaded regions presenting the 95% credible interval. The distribution of heterozygosity values used in each model is shown with a gray histogram, with the number of individuals shown on the right. Breeder Transition predictions are divided between locally hatched (E) and immigrant founder (F) individuals to separate effects of dispersal itself.

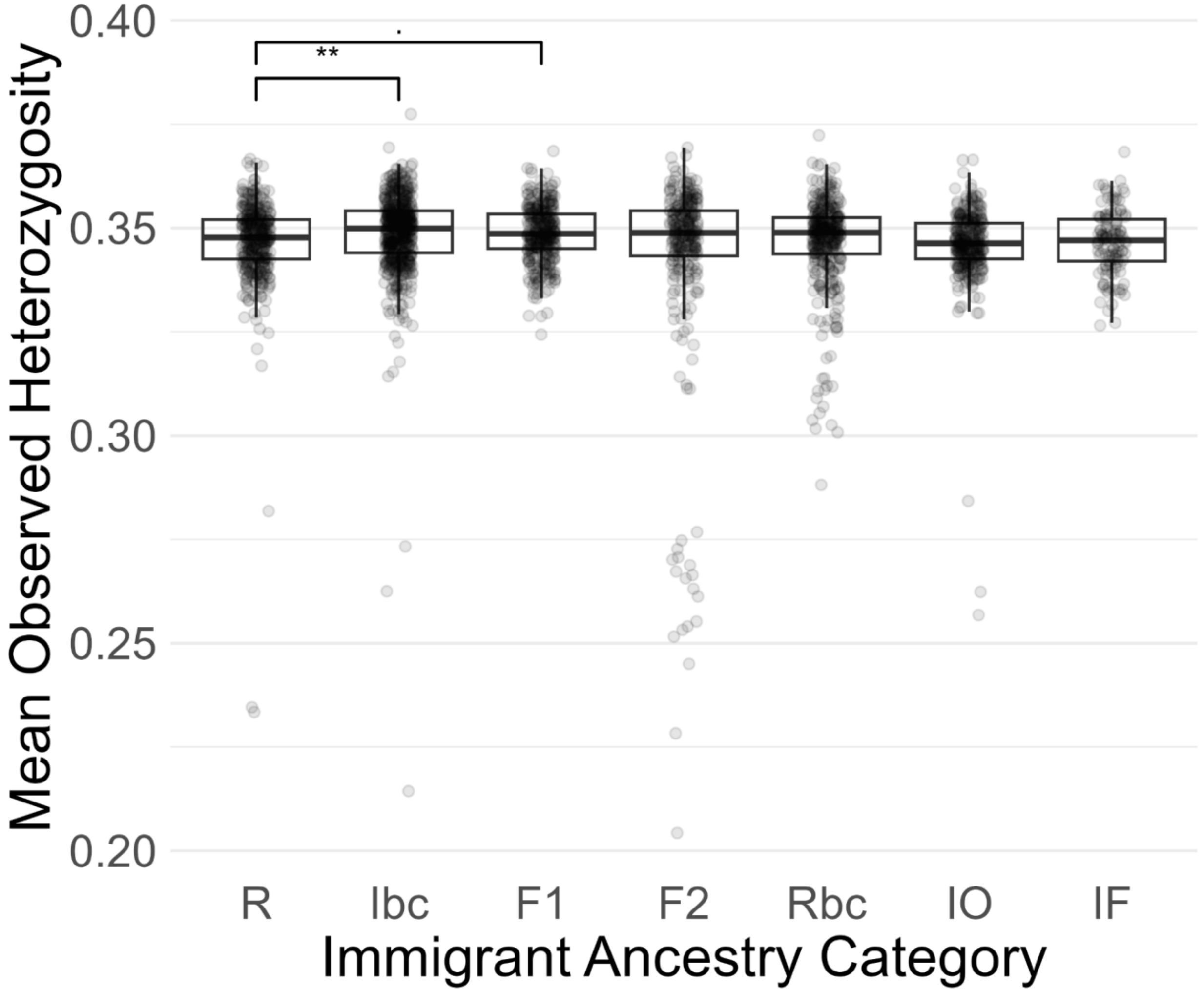


**Figure S11.** Mean observed heterozygosity for each immigrant ancestry category. Points represent individual values. Significance of Wilcoxon rank-sum test comparison between residents and other immigrant ancestry categories are shown, with no line indicating $p > 0.1$, . indicating $p < 0.1$, * indicating $p < 0.05$, and ** indicating $p < 0.01$.

| | Est. | | | 2.5 % | | | 97.5 % | | | Eff.Samp. | | |
|---|---|---|---|---|---|---|---|---|---|---|---|---|
| | (1) | (2) | (3) | (1) | (2) | (3) | (1) | (2) | (3) | (1) | (2) | (3) |
| Intercept | **1.566*** ** | **1.566*** ** | **1.574*** ** | 1.14 | 1.16 | 1.15 | 1.96 | 1.98 | 1.98 | 3750 | 3576 | 3750 |
| Resident Backcross | 0.258 | 0.260 | 0.253 | -0.26 | -0.28 | -0.28 | 0.81 | 0.81 | 0.77 | 3489 | 3573 | 3750 |
| F1 | **0.566*** | **0.571*** | **0.562*** | 0.06 | 0.03 | 0.08 | 1.05 | 1.02 | 1.06 | 3750 | 3750 | 3750 |
| F2 | 0.539 | 0.545 | 0.531 | -0.04 | -0.05 | -0.04 | 1.10 | 1.09 | 1.10 | 3750 | 3750 | 3750 |
| Immigrant Backcross | **0.565*** | **0.559*** | **0.553*** | 0.08 | 0.11 | 0.06 | 1.01 | 1.02 | 1.00 | 3750 | 3744 | 3750 |
| Immigrant Offspring | **0.558*** | **0.563*** | 0.550 | -0.00 | 0.01 | -0.03 | 1.08 | 1.12 | 1.09 | 4298 | 3517 | 3750 |
| Immigrant Founder | **0.460*** | **0.459*** | **0.450*** | 0.04 | 0.01 | 0.02 | 0.91 | 0.88 | 0.89 | 3528 | 3472 | 3750 |
| Sex | 0.296 | 0.296 | 0.285 | -0.22 | -0.24 | -0.25 | 0.82 | 0.83 | 0.81 | 3750 | 3984 | 3750 |
| Resident Backcross:Sex | 0.029 | 0.026 | 0.036 | -0.61 | -0.60 | -0.63 | 0.73 | 0.72 | 0.69 | 3750 | 3750 | 3750 |
| F1:Sex | -0.311 | -0.316 | -0.297 | -0.96 | -0.96 | -0.95 | 0.36 | 0.36 | 0.35 | 3750 | 4254 | 3750 |
| F2:Sex | -0.427 | -0.433 | -0.420 | -1.16 | -1.17 | -1.17 | 0.33 | 0.27 | 0.30 | 3750 | 3750 | 3750 |
| Immigrant Backcross:Sex | -0.426 | -0.415 | -0.407 | -1.05 | -1.03 | -1.03 | 0.16 | 0.19 | 0.20 | 3750 | 3759 | 3750 |
| Immigrant Offspring:Sex | -0.585 | -0.595 | -0.578 | -1.30 | -1.31 | -1.29 | 0.10 | 0.12 | 0.13 | 4008 | 3544 | 3750 |
| Immigrant Founder:Sex | -0.398 | -0.402 | -0.385 | -0.99 | -1.00 | -0.95 | 0.20 | 0.21 | 0.21 | 3750 | 4141 | 3750 |
| Year | 0.018 | 0.018 | 0.017 | 0.00 | 0.00 | 0.00 | 0.05 | 0.05 | 0.05 | 3750 | 3937 | 3750 |
| Year Interaction | 0.003 | 0.003 | 0.003 | -0.04 | -0.04 | -0.04 | 0.05 | 0.05 | 0.05 | 3750 | 3750 | 4076 |
| Parental ID | 0.077 | 0.077 | 0.075 | 0.00 | 0.00 | 0.00 | 0.18 | 0.18 | 0.17 | 3840 | 3750 | 3750 |
| Parental ID Interaction | 0.036 | 0.044 | 0.036 | -0.32 | -0.31 | -0.35 | 0.41 | 0.43 | 0.42 | 2771 | 3278 | 2415 |
| Intercept.B | **-3.181*** ** | **-3.165*** ** | **-3.315*** ** | -5.65 | -5.45 | -5.75 | -0.94 | -1.04 | -1.05 | 619 | 920 | 652 |
| Resident Backcross.B | -0.784 | -0.779 | -0.734 | -3.49 | -3.66 | -3.57 | 2.08 | 1.84 | 2.01 | 3576 | 3750 | 3750 |
| F1.B | 0.477 | 0.460 | 0.522 | -1.69 | -1.73 | -1.62 | 2.83 | 2.78 | 3.01 | 3502 | 3326 | 3261 |
| F2.B | 0.218 | 0.219 | 0.303 | -2.41 | -2.64 | -2.50 | 3.13 | 2.85 | 2.94 | 3049 | 3742 | 3032 |
| Immigrant Backcross.B | -0.839 | -0.851 | -0.823 | -3.17 | -3.09 | -3.36 | 1.56 | 1.47 | 1.36 | 3750 | 3729 | 3750 |

| | | | | | | | | | | | | |
|---|---|---|---|---|---|---|---|---|---|---|---|---|
| Immigrant Offspring.B | 0.850 | 0.846 | 0.925 | -1.79 | -1.65 | -1.55 | 3.31 | 3.29 | 3.46 | 3784 | 3507 | 2809 |
| Immigrant Founder.B | 0.025 | 0.021 | 0.073 | -2.04 | -1.96 | -1.87 | 2.06 | 2.02 | 2.23 | 3750 | 3979 | 3750 |
| Sex.B | 0.232 | 0.208 | 0.294 | -2.19 | -2.12 | -2.06 | 2.73 | 2.60 | 2.75 | 3750 | 3750 | 3124 |
| Resident Backcross:Sex.B | 0.455 | 0.510 | 0.397 | -3.25 | -2.65 | -3.15 | 3.55 | 3.89 | 3.67 | 3315 | 2775 | 2745 |
| F1:Sex.B | 0.120 | 0.163 | 0.090 | -2.90 | -2.67 | -2.68 | 2.93 | 3.03 | 3.13 | 3750 | 3750 | 3750 |
| F2:Sex.B | -1.410 | -1.352 | -1.529 | -5.42 | -4.93 | -5.85 | 2.49 | 2.73 | 2.09 | 1796 | 2744 | 1815 |
| Immigrant Backcross:Sex.B | 0.552 | 0.592 | 0.517 | -2.31 | -2.26 | -2.41 | 3.37 | 3.30 | 3.34 | 3750 | 3750 | 3656 |
| Immigrant Offspring:Sex.B | -1.200 | -1.189 | -1.269 | -4.48 | -4.36 | -4.58 | 1.98 | 2.03 | 1.91 | 3750 | 3750 | 3750 |
| Immigrant Founder:Sex.B | -0.130 | -0.079 | -0.190 | -3.03 | -2.63 | -2.91 | 2.51 | 2.67 | 2.67 | 3750 | 3750 | 2790 |
| Year.B | 0.128 | 0.125 | 0.133 | 0.00 | 0.00 | 0.00 | 0.49 | 0.48 | 0.51 | 891 | 1976 | 1172 |
| Year Interaction.B | 0.003 | 0.003 | 0.003 | -0.04 | -0.04 | -0.04 | 0.05 | 0.05 | 0.05 | 3750 | 3750 | 4076 |
| Parental ID.B | 3.245 | 3.060 | 3.616 | 0.00 | 0.00 | 0.00 | 9.89 | 9.07 | 11.15 | 179 | 324 | 195 |
| Parental ID Interaction.B | 0.036 | 0.044 | 0.036 | -0.32 | -0.31 | -0.35 | 0.41 | 0.43 | 0.42 | 2771 | 3278 | 2415 |
| DIC | 3924 | 3923 | 3921 | | | | | | | | | |
| N | 1406 | 1406 | 1406 | | | | | | | | | |

* p < 0.05, ** p < 0.01, *** p < 0.001

**Table S1.** Model results for breeder lifetime reproductive success. We present the mean posterior estimate (Est.), 95% credible interval, and effective sample size (Eff.Samp.) for each fixed and random effect across each of the 3 independent chains. Fixed effects with colons indicate interaction terms, with sex effects accounting for deviations in males relative to females. This model is divided into two sub-models, with the Poisson sub-model presented first, followed by the Binomial sub-model presented with .B appended to the effect names. Interaction random effects signify covariation between random effects of the two sub-models.

| | Est. | | | 2.5 % | | | 97.5 % | | | Eff.Samp. | | |
|---|---|---|---|---|---|---|---|---|---|---|---|---|
| | (1) | (2) | (3) | (1) | (2) | (3) | (1) | (2) | (3) | (1) | (2) | (3) |
| Intercept | **1.074**** | **1.086**** | **1.067**** | 0.34 | 0.32 | 0.35 | 1.82 | 1.74 | 1.74 | 3280 | 2918 | 2972 |
| Resident Backcross | 0.514 | 0.503 | 0.517 | -0.40 | -0.40 | -0.44 | 1.49 | 1.40 | 1.42 | 3750 | 3750 | 3750 |
| F1 | 0.751 | 0.742 | 0.775 | -0.11 | -0.08 | -0.11 | 1.63 | 1.60 | 1.61 | 2575 | 3753 | 3571 |
| F2 | 0.563 | 0.549 | 0.569 | -0.50 | -0.47 | -0.44 | 1.68 | 1.65 | 1.58 | 3750 | 3750 | 3750 |
| Immigrant Backcross | **0.913*** | **0.902*** | **0.922*** | 0.00 | 0.06 | 0.14 | 1.74 | 1.76 | 1.82 | 3750 | 3769 | 3579 |
| Immigrant Offspring | 0.761 | 0.749 | 0.765 | -0.23 | -0.28 | -0.30 | 1.88 | 1.76 | 1.76 | 3750 | 3750 | 3705 |
| Immigrant Founder | 0.562 | 0.554 | 0.572 | -0.18 | -0.16 | -0.12 | 1.33 | 1.26 | 1.29 | 3750 | 3750 | 3750 |
| Unknown Grandparents | -0.866 | -0.866 | -0.859 | -2.89 | -2.78 | -2.77 | 1.14 | 1.11 | 1.16 | 3420 | 3963 | 3750 |
| Sex | 0.742 | 0.733 | 0.748 | -0.11 | -0.12 | -0.03 | 1.65 | 1.56 | 1.62 | 3750 | 3400 | 3218 |
| Resident Backcross:Sex | -0.408 | -0.397 | -0.414 | -1.62 | -1.58 | -1.55 | 0.73 | 0.69 | 0.74 | 3750 | 3750 | 3468 |
| F1:Sex | -0.917 | -0.900 | -0.910 | -2.07 | -2.04 | -2.09 | 0.25 | 0.23 | 0.16 | 2513 | 3365 | 2935 |
| F2:Sex | -0.634 | -0.613 | -0.623 | -2.02 | -1.95 | -2.04 | 0.66 | 0.64 | 0.61 | 3750 | 3750 | 3750 |
| Immigrant Backcross:Sex | -1.002 | -0.994 | -1.004 | -2.14 | -2.12 | -2.18 | 0.07 | 0.07 | 0.02 | 3345 | 3327 | 2668 |
| Immigrant Offspring:Sex | -0.983 | -0.982 | -0.975 | -2.41 | -2.35 | -2.33 | 0.28 | 0.34 | 0.33 | 3482 | 3469 | 3218 |
| Immigrant Founder:Sex | -0.660 | -0.642 | -0.649 | -1.67 | -1.63 | -1.62 | 0.35 | 0.34 | 0.32 | 3750 | 3750 | 3460 |
| Unknown Grandparents:Sex | 2.640 | **2.643*** | **2.608*** | -0.20 | 0.07 | 0.04 | 5.27 | 5.49 | 5.36 | 3438 | 3750 | 3392 |
| Age | **-0.459**** | **-0.448**** | **-0.446**** | -0.89 | -0.92 | -0.90 | -0.05 | -0.06 | -0.05 | 1170 | 1086 | 1020 |
| $Age^2$ | **-0.290***** | **-0.291***** | **-0.291***** | -0.38 | -0.38 | -0.38 | -0.20 | -0.20 | -0.20 | 3750 | 3750 | 3750 |
| Year | 0.142 | 0.144 | 0.140 | 0.03 | 0.02 | 0.04 | 0.28 | 0.29 | 0.28 | 3564 | 3369 | 3460 |
| Parental ID | 0.589 | 0.577 | 0.575 | 0.00 | 0.00 | 0.00 | 1.20 | 1.18 | 1.18 | 2016 | 1975 | 1639 |
| Individual ID | 1.122 | 1.091 | 1.095 | 0.00 | 0.00 | 0.00 | 2.66 | 2.80 | 2.73 | 1142 | 1026 | 1032 |
| DIC | 3583 | 3586 | 3586 | | | | | | | | | |
| N | 3700 | 3700 | 3700 | | | | | | | | | |

* $p < 0.05$, ** $p < 0.01$, *** $p < 0.001$

**Table S2.** Model results for breeder survival. We present the mean posterior estimate (Est.), 95% credible interval, and effective sample size (Eff.Samp.) for each fixed and random effect across each of the 3 independent chains. Fixed effects with colons indicate interaction terms, with sex effects accounting for deviations in males relative to females.

| | Est. | | | 2.5 % | | | 97.5 % | | | Eff.Samp. | | |
|---|---|---|---|---|---|---|---|---|---|---|---|---|
| | (1) | (2) | (3) | (1) | (2) | (3) | (1) | (2) | (3) | (1) | (2) | (3) |
| Intercept | **0.979*** ** | **0.978*** ** | **0.979*** ** | 0.83 | 0.84 | 0.84 | 1.11 | 1.13 | 1.12 | 1305 | 1296 | 1180 |
| Resident Backcross | -0.021 | -0.018 | -0.019 | -0.21 | -0.21 | -0.21 | 0.18 | 0.18 | 0.17 | 1157 | 1232 | 1013 |
| F1 | -0.032 | -0.033 | -0.033 | -0.20 | -0.21 | -0.21 | 0.13 | 0.14 | 0.12 | 1330 | 1217 | 1107 |
| F2 | 0.002 | -0.003 | 0.001 | -0.21 | -0.21 | -0.19 | 0.19 | 0.20 | 0.21 | 1181 | 1226 | 1110 |
| Immigrant Backcross | 0.037 | 0.037 | 0.036 | -0.13 | -0.13 | -0.12 | 0.19 | 0.20 | 0.20 | 1284 | 1260 | 1152 |
| Immigrant Offspring | 0.033 | 0.031 | 0.034 | -0.15 | -0.16 | -0.15 | 0.23 | 0.22 | 0.23 | 1357 | 1250 | 1126 |
| Immigrant Founder | 0.002 | 0.002 | 0.002 | -0.13 | -0.15 | -0.14 | 0.15 | 0.16 | 0.15 | 1316 | 1220 | 1084 |
| Unknown Grandparents | -0.013 | -0.000 | -0.009 | -0.44 | -0.47 | -0.44 | 0.44 | 0.42 | 0.43 | 1444 | 1079 | 1129 |
| Sex | 0.011 | 0.014 | 0.011 | -0.16 | -0.16 | -0.17 | 0.18 | 0.19 | 0.17 | 1347 | 1114 | 1080 |
| Resident Backcross:Sex | 0.029 | 0.018 | 0.022 | -0.21 | -0.21 | -0.21 | 0.25 | 0.26 | 0.26 | 1270 | 1203 | 1065 |
| F1:Sex | 0.011 | 0.011 | 0.015 | -0.20 | -0.20 | -0.20 | 0.23 | 0.24 | 0.24 | 1319 | 1235 | 1052 |
| F2:Sex | -0.040 | -0.032 | -0.037 | -0.30 | -0.32 | -0.31 | 0.21 | 0.23 | 0.22 | 1248 | 1093 | 1038 |
| Immigrant Backcross:Sex | -0.050 | -0.055 | -0.052 | -0.26 | -0.27 | -0.25 | 0.15 | 0.15 | 0.15 | 1247 | 1211 | 1110 |
| Immigrant Offspring:Sex | -0.030 | -0.030 | -0.037 | -0.29 | -0.29 | -0.29 | 0.22 | 0.22 | 0.20 | 1282 | 1140 | 1084 |
| Immigrant Founder:Sex | -0.029 | -0.032 | -0.030 | -0.24 | -0.24 | -0.23 | 0.16 | 0.17 | 0.16 | 1234 | 1134 | 1076 |
| Unknown Grandparents:Sex | -0.005 | -0.019 | -0.007 | -0.53 | -0.54 | -0.51 | 0.54 | 0.51 | 0.56 | 1401 | 1119 | 1091 |
| Age | **0.093*** ** | **0.093*** ** | **0.093*** ** | 0.06 | 0.06 | 0.06 | 0.13 | 0.13 | 0.13 | 1099 | 951 | 1004 |
| $Age^2$ | **-0.044*** ** | **-0.045*** ** | **-0.044*** ** | -0.07 | -0.07 | -0.07 | -0.02 | -0.02 | -0.02 | 993 | 953 | 869 |
| Year | 0.009 | 0.009 | 0.009 | 0.00 | 0.00 | 0.00 | 0.02 | 0.02 | 0.02 | 1084 | 978 | 1026 |
| Year Interaction | -0.049 | -0.049 | -0.049 | -0.09 | -0.09 | -0.09 | -0.01 | -0.01 | -0.01 | 2616 | 2328 | 2017 |
| Parental ID | 0.001 | 0.000 | 0.000 | 0.00 | 0.00 | 0.00 | 0.00 | 0.00 | 0.00 | 1536 | 1422 | 1246 |
| Parental ID Interaction | -0.001 | -0.001 | -0.001 | -0.01 | -0.01 | -0.01 | 0.01 | 0.01 | 0.01 | 3485 | 3456 | 3435 |
| Individual ID | 0.001 | 0.001 | 0.001 | 0.00 | 0.00 | 0.00 | 0.00 | 0.00 | 0.00 | 1233 | 1219 | 1097 |
| Individual ID Interaction | -0.004 | -0.004 | -0.004 | -0.02 | -0.02 | -0.02 | 0.01 | 0.01 | 0.01 | 1593 | 1604 | 1618 |
| Intercept.B | **-1.251*** ** | **-1.245*** ** | **-1.246*** ** | -1.84 | -1.86 | -1.84 | -0.64 | -0.69 | -0.66 | 8000 | 8000 | 8000 |

| | | | | | | | | | | | | |
|---|---|---|---|---|---|---|---|---|---|---|---|---|
| Resident Backcross.B | 0.073 | 0.069 | 0.071 | -0.65 | -0.68 | -0.64 | 0.78 | 0.75 | 0.79 | 7387 | 8000 | 7042 |
| F1.B | -0.297 | -0.305 | -0.307 | -0.97 | -0.95 | -0.95 | 0.36 | 0.35 | 0.36 | 8000 | 8000 | 8000 |
| F2.B | -0.329 | -0.332 | -0.326 | -1.17 | -1.15 | -1.14 | 0.50 | 0.45 | 0.48 | 7732 | 8000 | 8000 |
| Immigrant Backcross.B | -0.255 | -0.255 | -0.257 | -0.91 | -0.89 | -0.90 | 0.39 | 0.40 | 0.41 | 8260 | 7713 | 7433 |
| Immigrant Offspring.B | -0.447 | -0.454 | -0.453 | -1.25 | -1.24 | -1.22 | 0.32 | 0.29 | 0.34 | 8000 | 7512 | 8000 |
| Immigrant Founder.B | 0.001 | -0.004 | -0.004 | -0.55 | -0.55 | -0.55 | 0.59 | 0.58 | 0.59 | 8000 | 8000 | 8000 |
| Unknown Grandparents.B | 0.522 | 0.490 | 0.481 | -1.14 | -1.25 | -1.24 | 2.22 | 2.12 | 2.12 | 8167 | 7510 | 8000 |
| Sex.B | -0.002 | -0.009 | -0.014 | -0.62 | -0.63 | -0.64 | 0.64 | 0.63 | 0.63 | 7497 | 8425 | 8000 |
| Resident Backcross:Sex.B | -0.429 | -0.424 | -0.418 | -1.29 | -1.27 | -1.29 | 0.42 | 0.48 | 0.43 | 6953 | 8000 | 8000 |
| F1:Sex.B | -0.033 | -0.021 | -0.014 | -0.86 | -0.90 | -0.84 | 0.81 | 0.79 | 0.81 | 7568 | 8000 | 8000 |
| F2:Sex.B | 0.253 | 0.257 | 0.262 | -0.79 | -0.76 | -0.79 | 1.23 | 1.23 | 1.23 | 7693 | 8000 | 8000 |
| Immigrant Backcross:Sex.B | 0.241 | 0.243 | 0.247 | -0.53 | -0.53 | -0.54 | 1.03 | 1.01 | 1.02 | 8000 | 8000 | 8459 |
| Immigrant Offspring:Sex.B | 0.563 | 0.570 | 0.573 | -0.46 | -0.41 | -0.42 | 1.50 | 1.55 | 1.50 | 8000 | 8000 | 8000 |
| Immigrant Founder:Sex.B | 0.081 | 0.088 | 0.092 | -0.68 | -0.64 | -0.65 | 0.83 | 0.84 | 0.81 | 8000 | 8000 | 8000 |
| Unknown Grandparents:Sex.B | -1.116 | -1.066 | -1.073 | -3.24 | -3.19 | -3.21 | 0.99 | 1.07 | 1.04 | 7464 | 7657 | 8000 |
| Age.B | **-0.471****** | **-0.470****** | **-0.470****** | -0.59 | -0.59 | -0.58 | -0.36 | -0.36 | -0.36 | 8000 | 8000 | 8000 |
| $Age^2$.B | **0.282****** | **0.283****** | **0.283****** | 0.20 | 0.21 | 0.20 | 0.36 | 0.36 | 0.36 | 8000 | 8593 | 8000 |
| Year.B | 0.562 | 0.561 | 0.562 | 0.27 | 0.27 | 0.28 | 0.91 | 0.91 | 0.93 | 8000 | 8000 | 8000 |
| Year Interaction.B | -0.049 | -0.049 | -0.049 | -0.09 | -0.09 | -0.09 | -0.01 | -0.01 | -0.01 | 2616 | 2328 | 2017 |
| Parental ID.B | 0.114 | 0.111 | 0.114 | 0.00 | 0.00 | 0.00 | 0.31 | 0.31 | 0.31 | 6224 | 6228 | 6462 |
| Parental ID Interaction.B | -0.001 | -0.001 | -0.001 | -0.01 | -0.01 | -0.01 | 0.01 | 0.01 | 0.01 | 3485 | 3456 | 3435 |
| Individual ID.B | 0.241 | 0.239 | 0.240 | 0.00 | 0.00 | 0.00 | 0.49 | 0.48 | 0.48 | 5330 | 4537 | 5505 |
| Individual ID Interaction.B | -0.004 | -0.004 | -0.004 | -0.02 | -0.02 | -0.02 | 0.01 | 0.01 | 0.01 | 1593 | 1604 | 1618 |
| DIC | 12332 | 12332 | 12331 | | | | | | | | | |
| N | 7400 | 7400 | 7400 | | | | | | | | | |

* p < 0.05, ** p < 0.01, *** p < 0.001

**Table S3.** Model results for breeder annual reproductive success. We present the mean posterior estimate (Est.), 95% credible interval, and effective sample size (Eff.Samp.) for each fixed and random effect across each of the 3 independent chains. Fixed effects with colons indicate interaction terms, with sex effects accounting for deviations in males relative to females. This model is divided into two sub-models, with the Poisson sub-model presented first, followed by the Binomial sub-model presented with .B appended to the effect names. Interaction random effects signify covariation between random effects of the two sub-models.

| | Est. | | | 2.5 % | | | 97.5 % | | | Eff.Samp. | | |
|---|---|---|---|---|---|---|---|---|---|---|---|---|
| | (1) | (2) | (3) | (1) | (2) | (3) | (1) | (2) | (3) | (1) | (2) | (3) |
| Intercept | -0.604 | -0.610 | -0.600 | -1.34 | -1.31 | -1.31 | 0.11 | 0.14 | 0.14 | 4000 | 4000 | 4000 |
| Resident Backcross | -0.453 | -0.447 | -0.455 | -1.26 | -1.24 | -1.19 | 0.33 | 0.31 | 0.40 | 3711 | 4000 | 4000 |
| F1 | -0.463 | -0.453 | -0.467 | -1.25 | -1.29 | -1.24 | 0.43 | 0.36 | 0.40 | 3800 | 4000 | 4000 |
| F2 | -0.431 | -0.424 | -0.429 | -1.30 | -1.31 | -1.29 | 0.43 | 0.41 | 0.43 | 4000 | 4000 | 4000 |
| Immigrant Backcross | -0.209 | -0.201 | -0.212 | -1.00 | -1.00 | -1.00 | 0.58 | 0.55 | 0.58 | 4000 | 4000 | 4000 |
| Immigrant Offspring | -0.702 | -0.689 | -0.702 | -1.56 | -1.60 | -1.65 | 0.15 | 0.17 | 0.12 | 4000 | 4000 | 4000 |
| Sex | -0.302 | -0.304 | -0.297 | -1.15 | -1.15 | -1.18 | 0.58 | 0.62 | 0.57 | 3813 | 4000 | 4000 |
| Resident Backcross:Sex | 0.451 | 0.458 | 0.444 | -0.54 | -0.56 | -0.50 | 1.37 | 1.38 | 1.48 | 3820 | 4000 | 4000 |
| F1:Sex | 0.152 | 0.151 | 0.146 | -0.89 | -0.83 | -0.86 | 1.18 | 1.16 | 1.18 | 3995 | 4000 | 4000 |
| F2:Sex | 0.588 | 0.586 | 0.583 | -0.42 | -0.49 | -0.47 | 1.69 | 1.62 | 1.67 | 4000 | 4022 | 4000 |
| Immigrant Backcross:Sex | 0.226 | 0.226 | 0.221 | -0.73 | -0.81 | -0.71 | 1.17 | 1.15 | 1.23 | 4000 | 4000 | 4000 |
| Immigrant Offspring:Sex | 0.409 | 0.398 | 0.403 | -0.68 | -0.70 | -0.67 | 1.46 | 1.47 | 1.48 | 4000 | 4000 | 4000 |
| Hatch Date | **-0.330*** ** | **-0.331*** ** | **-0.331*** ** | -0.47 | -0.45 | -0.46 | -0.20 | -0.19 | -0.19 | 4419 | 4000 | 4421 |
| Year | 0.194 | 0.193 | 0.196 | 0.04 | 0.03 | 0.03 | 0.41 | 0.41 | 0.41 | 4000 | 4000 | 4000 |
| Parental ID | 0.235 | 0.236 | 0.234 | 0.00 | 0.00 | 0.00 | 0.51 | 0.52 | 0.51 | 3531 | 3591 | 3165 |
| Nest ID | 1.429 | 1.411 | 1.418 | 0.91 | 0.87 | 0.88 | 1.99 | 1.96 | 1.98 | 3572 | 3731 | 3412 |
| DIC | 3613 | 3614 | 3613 | | | | | | | | | |
| N | 3020 | 3020 | 3020 | | | | | | | | | |

* p < 0.05, ** p < 0.01, *** p < 0.001

**Table S4.** Model results for juvenile survival. We present the mean posterior estimate (Est.), 95% credible interval, and effective sample size (Eff.Samp.) for each fixed and random effect across each of the 3 independent chains. Fixed effects with colons indicate interaction terms, with sex effects accounting for deviations in males relative to females.

| | Est. | | | 2.5 % | | | 97.5 % | | | Eff.Samp. | | |
|---|---|---|---|---|---|---|---|---|---|---|---|---|
| | (1) | (2) | (3) | (1) | (2) | (3) | (1) | (2) | (3) | (1) | (2) | (3) |
| Intercept | -0.636 | -0.631 | -0.637 | -1.33 | -1.36 | -1.34 | 0.06 | 0.03 | 0.05 | 6667 | 6667 | 6667 |
| Resident Backcross | 0.390 | 0.393 | 0.386 | -0.55 | -0.50 | -0.53 | 1.26 | 1.28 | 1.27 | 6667 | 6402 | 6392 |
| F1 | 0.335 | 0.333 | 0.337 | -0.51 | -0.51 | -0.45 | 1.18 | 1.15 | 1.26 | 6419 | 6667 | 6667 |
| F2 | 0.191 | 0.184 | 0.183 | -0.78 | -0.79 | -0.79 | 1.11 | 1.14 | 1.14 | 6667 | 6667 | 6667 |
| Immigrant Backcross | 0.489 | 0.480 | 0.483 | -0.32 | -0.37 | -0.35 | 1.32 | 1.29 | 1.28 | 6667 | 6667 | 6667 |
| Immigrant Offspring | 0.002 | 0.005 | 0.006 | -0.98 | -0.91 | -0.95 | 0.97 | 0.98 | 0.99 | 6667 | 6667 | 6667 |
| Immigrant Founder | **3.013*** ** | **3.007*** ** | **3.021*** ** | 1.94 | 1.89 | 1.89 | 4.15 | 4.14 | 4.17 | 6248 | 6667 | 6667 |
| Sex | **0.933*** | **0.923*** | 0.923 | -0.01 | -0.01 | -0.06 | 1.84 | 1.84 | 1.83 | 6667 | 6667 | 6667 |
| Resident Backcross:Sex | 0.006 | 0.007 | 0.018 | -1.15 | -1.25 | -1.18 | 1.26 | 1.16 | 1.32 | 6992 | 6667 | 6667 |
| F1:Sex | -0.472 | -0.460 | -0.465 | -1.69 | -1.59 | -1.65 | 0.67 | 0.69 | 0.66 | 6667 | 6124 | 6667 |
| F2:Sex | -0.060 | -0.043 | -0.043 | -1.36 | -1.33 | -1.37 | 1.29 | 1.27 | 1.31 | 6667 | 6909 | 6667 |
| Immigrant Backcross:Sex | -0.284 | -0.268 | -0.270 | -1.37 | -1.39 | -1.38 | 0.85 | 0.83 | 0.85 | 6667 | 6667 | 6667 |
| Immigrant Offspring:Sex | -0.080 | -0.071 | -0.070 | -1.36 | -1.34 | -1.39 | 1.28 | 1.20 | 1.27 | 7060 | 6720 | 6667 |
| Immigrant Founder:Sex | -0.302 | -0.296 | -0.291 | -1.64 | -1.59 | -1.56 | 1.02 | 1.04 | 1.08 | 6667 | 6973 | 6667 |
| Year | 0.064 | 0.065 | 0.064 | 0.00 | 0.00 | 0.00 | 0.19 | 0.19 | 0.19 | 6667 | 7995 | 6667 |
| Parental ID | 0.264 | 0.260 | 0.264 | 0.00 | 0.00 | 0.00 | 0.71 | 0.70 | 0.70 | 5879 | 5310 | 5557 |
| DIC | 1583 | 1583 | 1583 | | | | | | | | | |
| N | 1269 | 1269 | 1269 | | | | | | | | | |

* p < 0.05, ** p < 0.01, *** p < 0.001

**Table S5.** Model results for likelihood to become a breeder (breeder transition). We present the mean posterior estimate (Est.), 95% credible interval, and effective sample size (Eff.Samp.) for each fixed and random effect across each of the 3 independent chains. Fixed effects with colons indicate interaction terms, with sex effects accounting for deviations in males relative to females.

| | Est. | | | 2.5 % | | | 97.5 % | | | Eff.Samp. | | |
|---|---|---|---|---|---|---|---|---|---|---|---|---|
| | (1) | (2) | (3) | (1) | (2) | (3) | (1) | (2) | (3) | (1) | (2) | (3) |
| Intercept | **2.025***** | **2.026***** | **2.026***** | 1.91 | 1.91 | 1.91 | 2.14 | 2.14 | 2.14 | 3750 | 3750 | 3750 |
| Sex | -0.035 | -0.037 | -0.034 | -0.17 | -0.17 | -0.18 | 0.10 | 0.10 | 0.10 | 3750 | 3750 | 3750 |
| Heterozygosity | 0.013 | 0.012 | 0.012 | -0.11 | -0.12 | -0.11 | 0.14 | 0.15 | 0.14 | 3808 | 4026 | 3750 |
| Heterozygosity:Sex | -0.041 | -0.040 | -0.039 | -0.19 | -0.20 | -0.18 | 0.14 | 0.12 | 0.13 | 3999 | 3750 | 3936 |
| Year | 0.021 | 0.020 | 0.019 | 0.00 | 0.00 | 0.00 | 0.06 | 0.05 | 0.05 | 3155 | 3750 | 3750 |
| Year Interaction | 0.004 | 0.004 | 0.003 | -0.03 | -0.03 | -0.03 | 0.05 | 0.05 | 0.05 | 3750 | 3750 | 3750 |
| Parental ID | 0.065 | 0.065 | 0.065 | 0.00 | 0.00 | 0.00 | 0.16 | 0.15 | 0.15 | 3750 | 3750 | 3750 |
| Parental ID Interaction | 0.021 | 0.023 | 0.021 | -0.17 | -0.17 | -0.19 | 0.26 | 0.25 | 0.25 | 3750 | 3750 | 3574 |
| Intercept.B | **-2.676***** | **-2.670***** | **-2.696***** | -3.40 | -3.34 | -3.45 | -2.10 | -2.06 | -2.04 | 1326 | 1222 | 1070 |
| Sex.B | 0.082 | 0.083 | 0.080 | -0.47 | -0.48 | -0.47 | 0.66 | 0.64 | 0.66 | 3750 | 3750 | 3750 |
| Heterozygosity.B | -0.228 | -0.224 | -0.227 | -0.76 | -0.81 | -0.74 | 0.35 | 0.30 | 0.37 | 3175 | 3559 | 3514 |
| Heterozygosity:Sex.B | 0.209 | 0.204 | 0.199 | -0.50 | -0.45 | -0.45 | 0.89 | 0.95 | 0.91 | 3412 | 3750 | 3515 |
| Year.B | 0.079 | 0.078 | 0.075 | 0.00 | 0.00 | 0.00 | 0.30 | 0.30 | 0.29 | 3408 | 3396 | 3328 |
| Year Interaction.B | 0.004 | 0.004 | 0.003 | -0.03 | -0.03 | -0.03 | 0.05 | 0.05 | 0.05 | 3750 | 3750 | 3750 |
| Parental ID.B | 0.979 | 0.965 | 1.067 | 0.00 | 0.00 | 0.00 | 3.08 | 3.00 | 3.52 | 1014 | 992 | 914 |
| Parental ID Interaction.B | 0.021 | 0.023 | 0.021 | -0.17 | -0.17 | -0.19 | 0.26 | 0.25 | 0.25 | 3750 | 3750 | 3574 |
| DIC | 3925 | 3925 | 3924 | | | | | | | | | |
| N | 1406 | 1406 | 1406 | | | | | | | | | |

* p < 0.05, ** p < 0.01, *** p < 0.001

**Table S6.** Heterozygosity model results for breeder lifetime reproductive success. We present the mean posterior estimate (Est.), 95% credible interval, and effective sample size (Eff.Samp.) for each fixed and random effect across each of the 3 independent chains. Fixed effects with colons indicate interaction terms, with sex effects accounting for deviations in males relative to females. This model is divided into two sub-models, with the Poisson sub-model presented first, followed by the Binomial sub-model presented with .B appended to the effect names. Interaction random effects signify covariation between random effects of the two sub-models.

| | Est. | | | 2.5 % | | | 97.5 % | | | Eff.Samp. | | |
|---|---|---|---|---|---|---|---|---|---|---|---|---|
| | (1) | (2) | (3) | (1) | (2) | (3) | (1) | (2) | (3) | (1) | (2) | (3) |
| Intercept | **1.721*** ** | **1.718*** ** | **1.723*** ** | 1.48 | 1.47 | 1.47 | 1.97 | 1.97 | 1.97 | 3136 | 3080 | 3000 |
| Sex | 0.107 | 0.110 | 0.111 | -0.13 | -0.13 | -0.13 | 0.36 | 0.35 | 0.32 | 3263 | 3750 | 3750 |
| Age | **-0.278*** | **-0.275*** | **-0.269*** | -0.58 | -0.59 | -0.58 | -0.00 | -0.03 | -0.01 | 1891 | 2201 | 1723 |
| Age^2 | **-0.279*** ** | **-0.279*** ** | **-0.278*** ** | -0.37 | -0.37 | -0.36 | -0.20 | -0.20 | -0.20 | 3750 | 3750 | 4376 |
| Heterozygosity | 0.136 | 0.134 | 0.134 | -0.11 | -0.11 | -0.09 | 0.36 | 0.36 | 0.38 | 2825 | 3560 | 3750 |
| Heterozygosity:Sex | -0.089 | -0.086 | -0.086 | -0.38 | -0.36 | -0.38 | 0.20 | 0.20 | 0.19 | 3460 | 3564 | 3750 |
| Year | 0.125 | 0.122 | 0.122 | 0.03 | 0.02 | 0.03 | 0.25 | 0.24 | 0.25 | 3750 | 3506 | 3486 |
| Parental ID | 0.411 | 0.406 | 0.404 | 0.00 | 0.00 | 0.00 | 0.82 | 0.79 | 0.80 | 3144 | 3220 | 2908 |
| Individual ID | 0.489 | 0.482 | 0.463 | 0.00 | 0.00 | 0.00 | 1.42 | 1.37 | 1.33 | 1546 | 1822 | 1906 |
| DIC | 3635 | 3636 | 3637 | | | | | | | | | |
| N | 3700 | 3700 | 3700 | | | | | | | | | |

* p < 0.05, ** p < 0.01, *** p < 0.001

**Table S7.** Heterozygosity model results for breeder survival. We present the mean posterior estimate (Est.), 95% credible interval, and effective sample size (Eff.Samp.) for each fixed and random effect across each of the 3 independent chains. Fixed effects with colons indicate interaction terms, with sex effects accounting for deviations in males relative to females.

| | Est. | | | 2.5 % | | | 97.5 % | | | Eff.Samp. | | |
|---|---|---|---|---|---|---|---|---|---|---|---|---|
| | (1) | (2) | (3) | (1) | (2) | (3) | (1) | (2) | (3) | (1) | (2) | (3) |
| Intercept | **0.986*** ** | **0.985*** ** | **0.985*** ** | 0.93 | 0.93 | 0.93 | 1.04 | 1.04 | 1.04 | 1440 | 1650 | 1590 |
| Sex | -0.008 | -0.008 | -0.007 | -0.06 | -0.06 | -0.06 | 0.05 | 0.04 | 0.04 | 1085 | 1062 | 1146 |
| Age | **0.094*** ** | **0.094*** ** | **0.093*** ** | 0.06 | 0.06 | 0.06 | 0.13 | 0.13 | 0.13 | 1011 | 1086 | 1087 |
| Age^2 | **-0.044*** ** | **-0.044*** ** | **-0.043*** ** | -0.07 | -0.07 | -0.07 | -0.02 | -0.02 | -0.02 | 995 | 1073 | 1180 |
| Heterozygosity | -0.031 | -0.031 | -0.030 | -0.08 | -0.08 | -0.08 | 0.02 | 0.02 | 0.02 | 1186 | 1246 | 1233 |
| Heterozygosity:Sex | 0.026 | 0.026 | 0.026 | -0.04 | -0.04 | -0.04 | 0.09 | 0.09 | 0.08 | 1305 | 1283 | 1435 |
| Year | 0.009 | 0.009 | 0.009 | 0.00 | 0.00 | 0.00 | 0.02 | 0.02 | 0.02 | 1015 | 1038 | 1088 |
| Year Interaction | -0.048 | -0.049 | -0.048 | -0.09 | -0.09 | -0.09 | -0.01 | -0.01 | -0.01 | 2477 | 2036 | 2476 |
| Parental ID | 0.000 | 0.000 | 0.000 | 0.00 | 0.00 | 0.00 | 0.00 | 0.00 | 0.00 | 1274 | 1415 | 1383 |
| Parental ID Interaction | -0.001 | -0.001 | -0.001 | -0.01 | -0.01 | -0.01 | 0.01 | 0.01 | 0.01 | 3642 | 3359 | 3222 |
| Individual ID | 0.001 | 0.001 | 0.001 | 0.00 | 0.00 | 0.00 | 0.00 | 0.00 | 0.00 | 1058 | 775 | 1263 |
| Individual ID Interaction | -0.004 | -0.004 | -0.004 | -0.02 | -0.02 | -0.02 | 0.01 | 0.01 | 0.01 | 1581 | 1284 | 1863 |
| Intercept.B | **-1.352*** ** | **-1.352*** ** | **-1.351*** ** | -1.65 | -1.66 | -1.65 | -1.04 | -1.04 | -1.03 | 6950 | 8284 | 8563 |
| Sex.B | 0.032 | 0.030 | 0.032 | -0.17 | -0.17 | -0.16 | 0.22 | 0.23 | 0.23 | 8000 | 8000 | 8000 |
| Age.B | **-0.464*** ** | **-0.465*** ** | **-0.463*** ** | -0.57 | -0.58 | -0.58 | -0.34 | -0.35 | -0.35 | 7084 | 8000 | 8000 |
| Age^2.B | **0.277*** ** | **0.279*** ** | **0.277*** ** | 0.20 | 0.20 | 0.20 | 0.35 | 0.36 | 0.35 | 7714 | 8000 | 8000 |
| Heterozygosity.B | -0.173 | -0.173 | -0.174 | -0.38 | -0.37 | -0.37 | 0.03 | 0.03 | 0.03 | 8000 | 8602 | 8000 |
| Heterozygosity:Sex.B | **0.288* ** | **0.286* ** | **0.289* ** | 0.03 | 0.02 | 0.04 | 0.55 | 0.55 | 0.56 | 8718 | 8000 | 8000 |
| Year.B | 0.549 | 0.550 | 0.551 | 0.25 | 0.27 | 0.27 | 0.88 | 0.90 | 0.90 | 7329 | 8000 | 7709 |
| Year Interaction.B | -0.048 | -0.049 | -0.048 | -0.09 | -0.09 | -0.09 | -0.01 | -0.01 | -0.01 | 2477 | 2036 | 2476 |
| Parental ID.B | 0.113 | 0.111 | 0.113 | 0.00 | 0.00 | 0.00 | 0.30 | 0.30 | 0.30 | 6497 | 6350 | 6653 |
| Parental ID Interaction.B | -0.001 | -0.001 | -0.001 | -0.01 | -0.01 | -0.01 | 0.01 | 0.01 | 0.01 | 3642 | 3359 | 3222 |
| Individual ID.B | 0.216 | 0.220 | 0.218 | 0.00 | 0.00 | 0.00 | 0.45 | 0.45 | 0.45 | 6068 | 5489 | 5674 |
| Individual ID Interaction.B | -0.004 | -0.004 | -0.004 | -0.02 | -0.02 | -0.02 | 0.01 | 0.01 | 0.01 | 1581 | 1284 | 1863 |
| DIC | 12300 | 12300 | 12300 | | | | | | | | | |
| N | 7400 | 7400 | 7400 | | | | | | | | | |

* p < 0.05, ** p < 0.01, *** p < 0.001

**Table S8.** Heterozygosity model results for breeder annual reproductive success. We present the mean posterior estimate (Est.), 95% credible interval, and effective sample size (Eff.Samp.) for each fixed and random effect across each of the 3 independent chains. Fixed effects with colons indicate interaction terms, with sex effects accounting for deviations in males relative to females. This model is divided into two sub-models, with the Poisson sub-model presented first, followed by the Binomial sub-model presented with .B appended to the effect names. Interaction random effects signify covariation between random effects of the two sub-models.

| | Est. | | | 2.5 % | | | 97.5 % | | | Eff.Samp. | | |
|---|---|---|---|---|---|---|---|---|---|---|---|---|
| | (1) | (2) | (3) | (1) | (2) | (3) | (1) | (2) | (3) | (1) | (2) | (3) |
| Intercept | **-0.994*** ** | **-1.001*** ** | **-0.997*** ** | -1.23 | -1.26 | -1.24 | -0.74 | -0.76 | -0.74 | 4000 | 4000 | 4000 |
| Sex | 0.035 | 0.037 | 0.035 | -0.17 | -0.18 | -0.18 | 0.25 | 0.24 | 0.24 | 4003 | 4000 | 4000 |
| Hatch Date | **-0.336*** ** | **-0.335*** ** | **-0.334*** ** | -0.46 | -0.48 | -0.47 | -0.21 | -0.21 | -0.20 | 4000 | 4000 | 4000 |
| Heterozygosity | **0.401** ** | **0.400** ** | **0.396** ** | 0.15 | 0.13 | 0.14 | 0.68 | 0.67 | 0.65 | 4000 | 4000 | 4000 |
| Heterozygosity:Sex | **-0.392* ** | **-0.393* ** | **-0.386* ** | -0.72 | -0.73 | -0.71 | -0.09 | -0.08 | -0.08 | 4000 | 4000 | 4000 |
| Year | 0.182 | 0.183 | 0.184 | 0.02 | 0.03 | 0.03 | 0.37 | 0.39 | 0.38 | 3493 | 4000 | 3796 |
| Parental ID | 0.242 | 0.236 | 0.243 | 0.00 | 0.00 | 0.00 | 0.52 | 0.51 | 0.51 | 2976 | 3101 | 3147 |
| Nest ID | 1.318 | 1.342 | 1.332 | 0.80 | 0.81 | 0.85 | 1.85 | 1.90 | 1.94 | 3691 | 3587 | 3329 |
| DIC | 3607 | 3607 | 3606 | | | | | | | | | |
| N | 3020 | 3020 | 3020 | | | | | | | | | |

* p < 0.05, ** p < 0.01, *** p < 0.001

**Table S9.** Heterozygosity model results for juvenile survival. We present the mean posterior estimate (Est.), 95% credible interval, and effective sample size (Eff.Samp.) for each fixed and random effect across each of the 3 independent chains. Fixed effects with colons indicate interaction terms, with sex effects accounting for deviations in males relative to females.

| | Est. | | | 2.5 % | | | 97.5 % | | | Eff.Samp. | | |
|---|---|---|---|---|---|---|---|---|---|---|---|---|
| | (1) | (2) | (3) | (1) | (2) | (3) | (1) | (2) | (3) | (1) | (2) | (3) |
| Intercept | **2.266*** ** | **2.261*** ** | **2.263*** ** | 1.70 | 1.71 | 1.72 | 2.84 | 2.82 | 2.85 | 6667 | 7490 | 6667 |
| Sex | 0.360 | 0.356 | 0.355 | -0.44 | -0.45 | -0.45 | 1.16 | 1.16 | 1.15 | 6667 | 6667 | 6960 |
| Immigrant Founder | **-2.882*** ** | **-2.878*** ** | **-2.879*** ** | -3.51 | -3.50 | -3.49 | -2.27 | -2.30 | -2.28 | 6667 | 6667 | 6667 |
| Sex:Immigrant Founder | 0.553 | 0.556 | 0.553 | -0.35 | -0.34 | -0.35 | 1.39 | 1.42 | 1.42 | 6957 | 6667 | 7630 |
| Heterozygosity | 0.175 | 0.167 | 0.184 | -0.69 | -0.65 | -0.67 | 1.01 | 1.04 | 1.02 | 6667 | 6667 | 6667 |
| Heterozygosity:Sex | 0.253 | 0.246 | 0.232 | -1.31 | -1.42 | -1.30 | 1.72 | 1.75 | 1.77 | 6129 | 6826 | 6667 |
| Heterozygosity:Immigrant Founder | -0.155 | -0.144 | -0.164 | -1.04 | -1.03 | -1.01 | 0.78 | 0.78 | 0.82 | 6667 | 6667 | 6667 |
| Heterozygosity:Sex:Immigrant Founder | -0.068 | -0.064 | -0.049 | -1.65 | -1.70 | -1.62 | 1.50 | 1.55 | 1.57 | 6242 | 6574 | 6667 |
| Year | 0.058 | 0.059 | 0.062 | 0.00 | 0.00 | 0.00 | 0.17 | 0.18 | 0.19 | 6667 | 6667 | 6667 |
| Parental ID | 0.130 | 0.126 | 0.126 | 0.00 | 0.00 | 0.00 | 0.43 | 0.42 | 0.42 | 6272 | 6407 | 6375 |
| DIC | 1514 | 1514 | 1514 | | | | | | | | | |
| N | 1269 | 1269 | 1269 | | | | | | | | | |

* p < 0.05, ** p < 0.01, *** p < 0.001

**Table S10.** Heterozygosity model results for likelihood to become a breeder (breeder transition). We present the mean posterior estimate (Est.), 95% credible interval, and effective sample size (Eff.Samp.) for each fixed and random effect across each of the 3 independent chains. Fixed effects with colons indicate interaction terms, with sex effects accounting for deviations in males relative to females.

| | Breeder Lifetime Reproductive Success | |
|---|---|---|
| | Female | Male |
| Resident | 4.61 (2.9, 6.91)<br>-3.22 (-6.06, -1.28)<br>1.57 (1.15, 1.98) | 6.09 (4.06, 8.74)<br>-2.98 (-5.32, -1.32)<br>1.86 (1.5, 2.22) |
| Resident Backcross | 6.11 (4.27, 8.41)<br>-3.99 (-6.57, -2.1)<br>1.83 (1.49, 2.16) | 8.22 (6.21, 10.59)<br>-3.29 (-5.64, -1.84)<br>2.15 (1.89, 2.41) |
| F1 | 7.89 (5.77, 10.49)<br>-2.73 (-4.74, -1.42)<br>2.14 (1.86, 2.41) | 7.55 (5.39, 10.22)<br>-2.37 (-4.11, -1.15)<br>2.12 (1.82, 2.41) |
| F2 | 7.79 (5.03, 11.38)<br>-2.97 (-5.48, -1.21)<br>2.11 (1.72, 2.49) | 7.09 (4.93, 9.83)<br>-4.16 (-7.41, -2.07)<br>1.97 (1.63, 2.31) |
| Immigrant Backcross | 8.23 (6.51, 10.3)<br>-4.06 (-6.75, -2.47)<br>2.13 (1.9, 2.36) | 7.11 (5.57, 8.91)<br>-3.26 (-5.31, -1.98)<br>2 (1.77, 2.23) |
| Immigrant Offspring | 7.56 (4.8, 10.99)<br>-2.35 (-4.57, -0.81)<br>2.13 (1.75, 2.51) | 6.02 (4.19, 8.34)<br>-3.32 (-5.56, -1.66)<br>1.83 (1.5, 2.16) |
| Immigrant Founder | 7.23 (6.07, 8.52)<br>-3.18 (-5.17, -2.1)<br>2.02 (1.86, 2.18) | 6.48 (5.07, 8.19)<br>-3.07 (-5.36, -1.8)<br>1.92 (1.7, 2.15) |

**Table S11.** Posterior mean estimates for breeder lifetime reproductive success. Back-transformed values are presented on top, followed by the binomial sub-model latent values in the middle, and Poisson sub-model latent values on the bottom, with the 95% credible interval in parentheses.

| | Breeder Lifetime Reproductive Success | | |
|---|---|---|---|
| | Female | Male | Sex Difference |
| Rbc | 45% | 85% | 80% |
| VS | 18% | 23% | 61% |
| μ(F1, R) | 45% | 82% | 75% |
| F1 | **90%** | 86% | 44% |
| VS | 52% | 82% | 74% |
| μ(R, IO) | **93%** | **93%** | 48% |
| F2 | 66% | 57% | 41% |
| VS | 42% | 11% | 23% |
| μ(F1, μ(R, IO)) | 70% | 48% | 33% |
| Ibc | 65% | 61% | 45% |
| VS | 3% | 31% | 85% |
| μ(F1, IO) | 49% | 57% | 55% |
| IO VS R | **95%** | 49% | **9%** |
| | 76% | 38% | 22% |
| | 98% | 46% | **5%** |
| IO VS IF | 56% | 35% | 36% |
| | 83% | 39% | 19% |
| | 69% | 33% | 25% |

**Table S12.** Proportion of the posterior distribution of comparisons between ancestry categories greater than zero for breeder lifetime reproductive success. For sex differences, values greater than zero indicate that males exceed the expected phenotype more than females. Back-transformed values are presented on top, followed by the binomial sub-model latent values in the middle, and Poisson sub-model latent values on the bottom.

| | Lifetime Reproductive Success (Binomial) | |
|---|---|---|
| CGE | Female | Male |
| Additive | -0.07 (-2.59, 1.99)<br>5% | 0.03 (-1.65, 1.43)<br>1% |
| Dominance | -2.07 (-6.82, 1.67)<br>64% | 2.16 (-2.13, 6.39)<br>65% |
| Add. X Add.<br>Epistasis | -4.19 (-13.59, 3.3)<br>65% | 3.54 (-4.91, 11.9)<br>55% |
| Add. X Dom.<br>Epistasis | 2.31 (-0.81, 5.11)<br>85% | -0.37 (-3.4, 1.98)<br>19% |
| Dom. X Dom.<br>Epistasis | -0.51 (-3.44, 1.8)<br>28% | 0.2 (-1.86, 1.93)<br>15% |

**Table S13.** Posterior means of the composite genetic element (CGE) estimates for the binomial sub-model of breeder lifetime reproductive success. The 95% credible interval is presented in parentheses, with the credible interval that does not overlap zero presented below.

| | Lifetime Reproductive Success (Poisson) | |
|---|---|---|
| CGE | Female | Male |
| Additive | 0.3 (-0.09, 0.63)<br>86% | -0.14 (-0.48, 0.13)<br>60% |
| Dominance | -0.12 (-0.99, 0.62)<br>21% | 0.34 (-0.42, 0.98)<br>62% |
| Add. X Add. Epistasis | -0.52 (-2.22, 0.9)<br>45% | 0.41 (-1.06, 1.66)<br>41% |
| Add. X Dom. Epistasis | 0.15 (-0.44, 0.63)<br>37% | -0.2 (-0.7, 0.23)<br>55% |
| Dom. X Dom. Epistasis | 0.02 (-0.46, 0.43)<br>7% | -0.13 (-0.55, 0.21)<br>46% |

**Table S14.** Posterior means of the composite genetic element (CGE) estimates for the Poisson sub-model of breeder lifetime reproductive success. The 95% credible interval is presented in parentheses, with the credible interval that does not overlap zero presented below.

| | Breeder Survival | |
|---|---|---|
| | Female | Male |
| Resident | 0.74 (0.58, 0.85)<br>1.08 (0.32, 1.76) | 0.86 (0.77, 0.92)<br>1.82 (1.22, 2.41) |
| Resident Backcross | 0.83 (0.71, 0.9)<br>1.59 (0.91, 2.23) | 0.87 (0.79, 0.92)<br>1.92 (1.35, 2.47) |
| F1 | 0.86 (0.77, 0.92)<br>1.83 (1.21, 2.41) | 0.84 (0.73, 0.91)<br>1.66 (0.98, 2.26) |
| F2 | 0.83 (0.67, 0.92)<br>1.64 (0.73, 2.46) | 0.85 (0.73, 0.92)<br>1.75 (0.98, 2.48) |
| Immigrant Backcross | 0.88 (0.81, 0.92)<br>1.99 (1.44, 2.51) | 0.85 (0.76, 0.9)<br>1.73 (1.16, 2.23) |
| Immigrant Offspring | 0.86 (0.73, 0.93)<br>1.83 (1, 2.63) | 0.82 (0.69, 0.91)<br>1.6 (0.81, 2.32) |
| Immigrant Founder | 0.84 (0.77, 0.88)<br>1.64 (1.21, 2) | 0.85 (0.77, 0.9)<br>1.73 (1.19, 2.21) |

**Table S15.** Posterior mean estimates for breeder survival. Back-transformed values are presented above the latent values, with the 95% credibility interval in parentheses.

| | Breeder Survival | | |
|---|---|---|---|
| | Female | Male | Sex Difference |
| Rbc | 69% | 72% | 47% |
| VS | 64% | 71% | 54% |
| μ(F1, R) | | | |
| F1 | 88% | 49% | 17% |
| VS | 84% | 47% | 21% |
| μ(R, IO) | | | |
| F2 | 55% | 58% | 52% |
| VS | 49% | 56% | 55% |
| μ(F1, μ(R, IO)) | | | |
| Ibc | 69% | 64% | 47% |
| VS | 67% | 63% | 45% |
| μ(F1, IO) | | | |
| IO VS R | **93%** | 32% | **6%** |
| | **93%** | 32% | **7%** |
| IO VS IF | 68% | 37% | 29% |
| | 68% | 37% | 28% |

**Table S16.** Proportion of the posterior distribution of comparisons between ancestry categories greater than zero for breeder survival. For sex differences, values greater than zero indicate that males exceed the expected phenotype more than females. Back-transformed values are presented above the latent values.

| | Breeder Survival | |
| --- | --- | --- |
| CGE | Female | Male |
| Additive | 0.4 (-0.39, 1.08)<br>68% | -0.19 (-0.92, 0.38)<br>40% |
| Dominance | 0.49 (-1.32, 2.06)<br>39% | 0.12 (-1.54, 1.47)<br>13% |
| Add. X Add.<br>Epistasis | 0.61 (-2.96, 3.68)<br>25% | 0.29 (-2.91, 2.91)<br>14% |
| Add. X Dom.<br>Epistasis | -0.3 (-1.53, 0.68)<br>36% | -0.21 (-1.26, 0.66)<br>31% |
| Dom. X Dom.<br>Epistasis | 0.02 (-0.93, 0.81)<br>5% | -0.08 (-0.92, 0.62)<br>15% |

**Table S17.** Posterior means of the composite genetic element (CGE) estimates for breeder survival. The 95% credible interval is presented in parentheses, with the credible interval that does not overlap zero presented below.

| | Annual Reproductive Success | |
|---|---|---|
| | Female | Male |
| Resident | 2.06 (1.65, 2.49) | 2.09 (1.75, 2.44) |
| | -1.25 (-1.84, -0.66) | -1.26 (-1.75, -0.77) |
| | 0.98 (0.83, 1.12) | 0.99 (0.88, 1.1) |
| Resident Backcross | 1.99 (1.61, 2.39) | 2.25 (1.94, 2.56) |
| | -1.18 (-1.72, -0.65) | -1.61 (-2.09, -1.14) |
| | 0.96 (0.82, 1.09) | 0.99 (0.89, 1.09) |
| F1 | 2.12 (1.82, 2.43) | 2.19 (1.87, 2.5) |
| | -1.55 (-2.03, -1.07) | -1.58 (-2.08, -1.09) |
| | 0.95 (0.84, 1.05) | 0.97 (0.86, 1.08) |
| F2 | 2.2 (1.77, 2.65) | 2.05 (1.65, 2.45) |
| | -1.58 (-2.24, -0.92) | -1.33 (-1.92, -0.75) |
| | 0.98 (0.82, 1.13) | 0.95 (0.81, 1.09) |
| Immigrant Backcross | 2.25 (1.95, 2.56) | 2.07 (1.77, 2.36) |
| | -1.5 (-1.96, -1.05) | -1.27 (-1.7, -0.85) |
| | 1.02 (0.92, 1.11) | 0.98 (0.88, 1.07) |
| Immigrant Offspring | 2.32 (1.92, 2.74) | 2.04 (1.63, 2.46) |
| | -1.7 (-2.34, -1.07) | -1.14 (-1.71, -0.57) |
| | 1.01 (0.87, 1.15) | 0.99 (0.85, 1.13) |
| Immigrant Founder | 2.07 (1.82, 2.32) | 2 (1.7, 2.29) |
| | -1.25 (-1.6, -0.9) | -1.17 (-1.59, -0.75) |
| | 0.98 (0.91, 1.05) | 0.96 (0.87, 1.06) |

**Table S18.** Posterior mean estimates for annual reproductive success. Back-transformed values are presented on top, followed by the binomial sub-model latent values in the middle, and Poisson sub-model latent values on the bottom, with the 95% credibility interval in parentheses.

| | Annual Reproductive Success | | |
|---|---|---|---|
| | Female | Male | Sex Difference |
| Rbc | 32% | 74% | 79% |
| VS | 78% | 22% | 13% |
| μ(F1, R) | 49% | 59% | 57% |
| F1 | 36% | 75% | 76% |
| VS | 39% | **7%** | 21% |
| μ(R, IO) | 25% | 38% | 61% |
| F2 | 57% | 35% | 35% |
| VS | 43% | 58% | 61% |
| μ(F1, μ(R, IO)) | 54% | 37% | 38% |
| Ibc | 58% | 39% | 37% |
| VS | 69% | 66% | 46% |
| μ(F1, IO) | 72% | 46% | 31% |
| IO VS R | 83% | 41% | 19% |
| | 13% | 64% | 88% |
| | 63% | 50% | 41% |
| IO VS IF | 88% | 57% | 25% |
| | **7%** | 54% | 87% |
| | 66% | 64% | 50% |

**Table S19.** Proportion of the posterior distribution of comparisons between ancestry categories greater than zero for annual reproductive success. For sex differences, values greater than zero indicate that males exceed the expected phenotype more than females. Back-transformed values are presented on top, followed by the binomial sub-model latent values in the middle, and Poisson sub-model latent values on the bottom.

| | Annual Reproductive Success (Binomial) | |
|---|---|---|
| CGE | Female | Male |
| Additive | -0.33 (-0.9, 0.15)<br>74% | 0.34 (-0.15, 0.75)<br>83% |
| Dominance | 0.43 (-0.93, 1.58)<br>47% | -0.41 (-1.57, 0.57)<br>52% |
| Add. X Add. Epistasis | 0.95 (-1.71, 3.19)<br>51% | -0.44 (-2.7, 1.48)<br>30% |
| Add. X Dom. Epistasis | -0.41 (-1.29, 0.33)<br>64% | 0.16 (-0.6, 0.79)<br>32% |
| Dom. X Dom. Epistasis | -0.1 (-0.79, 0.48)<br>23% | 0.28 (-0.31, 0.78)<br>66% |

**Table S20.** Posterior means of the composite genetic element (CGE) estimates for the binomial sub-model of breeder annual reproductive success. The 95% credible interval is presented in parentheses, with the credible interval that does not overlap zero presented below.

| | Annual Reproductive Success (Poisson) | |
|---|---|---|
| CGE | Female | Male |
| Additive | 0.06 (-0.1, 0.19)<br>52% | -0.02 (-0.14, 0.08)<br>24% |
| Dominance | -0.01 (-0.34, 0.28)<br>4% | 0.05 (-0.25, 0.31)<br>25% |
| Add. X Add. Epistasis | 0.03 (-0.62, 0.6)<br>7% | 0.12 (-0.47, 0.63)<br>31% |
| Add. X Dom. Epistasis | -0.03 (-0.25, 0.16)<br>17% | -0.03 (-0.23, 0.13)<br>27% |
| Dom. X Dom. Epistasis | 0.04 (-0.14, 0.19)<br>33% | -0.02 (-0.17, 0.11)<br>19% |

**Table S21.** Posterior means of the composite genetic element (CGE) estimates for the Poisson sub-model of breeder annual reproductive success. The 95% credible interval is presented in parentheses, with the credible interval that does not overlap zero presented below.

| | Juvenile Survival | |
|---|---|---|
| | Female | Male |
| Resident | 0.36 (0.21, 0.53)<br>-0.6 (-1.35, 0.11) | 0.29 (0.16, 0.45)<br>-0.91 (-1.63, -0.2) |
| Resident Backcross | 0.26 (0.19, 0.34)<br>-1.06 (-1.46, -0.66) | 0.29 (0.21, 0.37)<br>-0.91 (-1.3, -0.53) |
| F1 | 0.26 (0.18, 0.35)<br>-1.07 (-1.52, -0.63) | 0.23 (0.16, 0.32)<br>-1.22 (-1.67, -0.77) |
| F2 | 0.27 (0.18, 0.37)<br>-1.03 (-1.54, -0.53) | 0.32 (0.22, 0.44)<br>-0.75 (-1.27, -0.24) |
| Immigrant Backcross | 0.31 (0.23, 0.4)<br>-0.81 (-1.2, -0.43) | 0.29 (0.22, 0.38)<br>-0.89 (-1.28, -0.5) |
| Immigrant Offspring | 0.22 (0.14, 0.32)<br>-1.3 (-1.85, -0.76) | 0.24 (0.15, 0.34)<br>-1.2 (-1.75, -0.67) |

**Table S22.** Posterior mean estimates for Juvenile survival. Back-transformed values are presented above the latent values, with the 95% credibility interval in parentheses.

| | Juvenile Survival | | |
|---|---|---|---|
| | Female | Male | Sex Difference |
| Rbc | 19% | 70% | 87% |
| VS | 21% | 72% | 87% |
| μ(F1, R) | | | |
| F1 | 32% | 27% | 47% |
| VS | 35% | 29% | 44% |
| μ(R, IO) | | | |
| F2 | 44% | **90%** | 88% |
| VS | 47% | **91%** | 87% |
| μ(F1, μ(R, IO)) | | | |
| Ibc | **93%** | 90% | 43% |
| VS | **94%** | 91% | 43% |
| μ(F1, IO) | | | |
| IO VS R | **6%** | 25% | 78% |
| | **6%** | 25% | 77% |

**Table S23.** Proportion of the posterior distribution of comparisons between ancestry categories greater than zero for juvenile survival. For sex differences, values greater than zero indicate that males exceed the expected phenotype more than females. Back-transformed values are presented above the latent values.

| | Juvenile Survival | |
|---|---|---|
| CGE | Female | Male |
| Additive | 0.24 (-0.25, 0.66)<br>65% | 0.02 (-0.48, 0.42)<br>7% |
| Dominance | 0.14 (-0.96, 1.08)<br>20% | -0.38 (-1.5, 0.57)<br>49% |
| Add. X Add. Epistasis | 0.39 (-1.74, 2.2)<br>28% | -0.6 (-2.75, 1.24)<br>42% |
| Add. X Dom. Epistasis | -0.17 (-0.91, 0.44)<br>36% | -0.09 (-0.83, 0.54)<br>18% |
| Dom. X Dom. Epistasis | 0.59 (-0.07, 1.15)<br>92% | 0.16 (-0.48, 0.7)<br>39% |

**Table S24.** Posterior means of the composite genetic element (CGE) estimates for juvenile survival. The 95% credible interval is presented in parentheses, with the credible interval that does not overlap zero presented below.

| | Breeder Transition | |
|---|---|---|
| | Female | Male |
| Resident | 0.35 (0.21, 0.51)<br>-0.63 (-1.34, 0.05) | 0.57 (0.4, 0.73)<br>0.29 (-0.4, 1) |
| Resident Backcross | 0.44 (0.3, 0.59)<br>-0.24 (-0.84, 0.35) | 0.66 (0.53, 0.78)<br>0.69 (0.12, 1.28) |
| F1 | 0.43 (0.31, 0.55)<br>-0.3 (-0.8, 0.2) | 0.54 (0.41, 0.67)<br>0.16 (-0.37, 0.7) |
| F2 | 0.39 (0.25, 0.55)<br>-0.45 (-1.12, 0.21) | 0.6 (0.43, 0.76)<br>0.43 (-0.27, 1.16) |
| Immigrant Backcross | 0.46 (0.36, 0.57)<br>-0.15 (-0.59, 0.29) | 0.62 (0.51, 0.73)<br>0.5 (0.03, 0.99) |
| Immigrant Offspring | 0.35 (0.21, 0.51)<br>-0.63 (-1.31, 0.03) | 0.55 (0.39, 0.71)<br>0.22 (-0.44, 0.89) |
| Immigrant Founder | 0.91 (0.83, 0.97)<br>2.38 (1.61, 3.41) | 0.95 (0.88, 0.99)<br>3.01 (2.02, 4.22) |

**Table S25.** Posterior mean estimates for likelihood to become a breeder (breeder transition). Back-transformed values are presented above the latent values, with the 95% credibility interval in parentheses.

| | Breeder Transition | | |
|---|---|---|---|
| | Female | Male | Sex Difference |
| Rbc | 72% | **90%** | 69% |
| VS | 73% | **90%** | 68% |
| μ(F1, R) | | | |
| F1 | 83% | 40% | 20% |
| VS | 84% | 40% | 19% |
| μ(R, IO) | | | |
| F2 | 51% | 71% | 65% |
| VS | 52% | 70% | 65% |
| μ(F1, μ(R, IO)) | | | |
| Ibc | 85% | 84% | 50% |
| VS | 85% | 83% | 50% |
| μ(F1, IO) | | | |
| IO VS R | 50% | 45% | 46% |
| | 50% | 45% | 46% |
| IO VS I | **0%** | **0%** | 93% |
| | **0%** | **0%** | 64% |

**Table S26.** Proportion of the posterior distribution of comparisons between ancestry categories greater than zero for likelihood to become a breeder (breeder transition). For sex differences, values greater than zero indicate that males exceed the expected phenotype more than females. Back-transformed values are presented above the latent values.

| | Breeder Transition | |
|---|---|---|
| CGE | Female | Male |
| Additive | 0.09 (-0.61, 0.7)<br>20% | -0.19 (-0.93, 0.43)<br>39% |
| Dominance | 0.67 (-0.85, 1.96)<br>61% | 0.29 (-1.34, 1.63)<br>27% |
| Add. X Add. Epistasis | 1 (-1.97, 3.54)<br>50% | 0.67 (-2.5, 3.3)<br>32% |
| Add. X Dom. Epistasis | -0.52 (-1.58, 0.33)<br>68% | -0.56 (-1.63, 0.34)<br>70% |
| Dom. X Dom. Epistasis | 0.09 (-0.76, 0.81)<br>16% | -0.16 (-1.03, 0.58)<br>28% |

**Table S27.** Posterior means of the composite genetic element (CGE) estimates for the likelihood to become a breeder (breeder transition). The 95% credible interval is presented in parentheses, with the credible interval that does not overlap zero presented below
